# Characterizing the Responses of Camelina and Sorghum to Environmental Stress through a Multi-Modal Approach

**DOI:** 10.1101/2024.10.30.621092

**Authors:** Kati A. Seitz, Demosthenes P. Morales, Steven T. Bristow, Erica G. Pauer, Duncan P. Ryan, Raymond T. Newell, David T. Hanson, James H. Werner

## Abstract

Due to their sessile nature, plants are unable to escape environmental factors that negatively impact health, resulting in losses to agricultural productivity. Rapid, non-invasive tools to detect plant stress response are essential for optimizing resource efficiency and mitigating the effects of extreme environmental pressures. However, many existing methods are either invasive, incompatible with other measurement techniques, or have not been applied to a wide range of varying environmental factors. In this study, we assess the physiological responses of four week old camelina (*Camelina sativa*) and sorghum (*Sorghum bicolor*) to chitosan, cold, drought, and both acute and chronic salt stress. Several plant characteristics were measured in parallel during stress exposure, including fluorescence and gas exchange parameters (MultispeQ and LI-6800), tissue electrical impedance with wearable biosensors (Multi-PIP), and biochemical properties via Fourier-transform infrared (FTIR) spectroscopy. We compiled unique profiles for whole plant physiological changes in response to environmental stress, demonstrating that certain aspects of plant health and makeup underwent alterations on differing temporal scales. This finding emphasizes the need for a comprehensive multi-modal approach to rapidly and accurately perform remote sensing of plant health in the field. Physiological parameters such as leaf impedance were also observed to rapidly change in response to treatment and can be leveraged to detect very early signs of plant perturbation. This research establishes the utility of a holistic phenotyping approach to inform agricultural strategies aimed at enhancing crop resilience under changing environmental conditions.

## Introduction

Abiotic stressors such as drought, salinity, and extreme temperatures are major limiting factors for plant growth and agricultural productivity globally, a challenge that will only intensify with the progression of climate change (Zhu, 2016). These environmental conditions negatively impact physiological processes, ultimately reducing crop yields and increasing plant mortality (Valliyodan and Nguyen, 2006). Therefore, the detection and management of environmentally-induced plant stress is critical to improving agricultural productivity and sustainability (Boyer, 1982; Mittler, 2006). However, traditional methods of measuring plant stress, such as biomass analysis or biochemical assays, are often time-consuming, invasive, and provide limited insight into dynamic physiological processes (Araus and Cairns, 2014), making them ill-suited for field-based monitoring.

Real-time, non-destructive quantification of plant stress is becoming increasingly attainable with the rise of smart agriculture—a farming approach that leverages sensor technologies, data analytics, and automation to optimize resource use and enhance crop productivity. This has resulted in a range of plant phenotyping methods being developed for rapid, continuous and accurate assessment of plant properties related to health and yield in the field (Jones and Vaughan, 2010; Zarco-Tejada et al., 2013; Ray et al., 2017). These technologies allow farmers to make informed decisions about irrigation, fertilization, and pest management. However, many of these tools either capture plant stress indirectly via soil or atmosphere parameters, which are susceptible to environmental influences, lack the sensitivity to detect early stages or different types of stress response, or fail to complement other methods, resulting in incomplete assessments of plant physiology (Chaves et al., 2009). Therefore, rigorous analysis of data compatibility and applicability for both pre-existing and novel tools is critical for assembling phenotyping platforms to reduce the negative impacts of environmental stress on plant growth and yield. Advancements in this field are also crucial for developing rigorous assessment and selection methodologies for resilient crop species, particularly those capable of thriving under diverse environmental conditions while maintaining productivity.

Camelina (*Camelina sativa*) and sorghum (*Sorghum bicolor*) are two crop species of emerging importance for biofuel production and global food security with relevance to both traditional and novel cropping systems (Johnson and Kershen, 2013; Li et al., 2019). Camelina, a C_3_ oilseed crop, is frost-tolerant and possesses a high resistance to drought, granting adaptability to a wide range of environments (Moser et al., 2010; Obour et al., 2015; Berti et al., 2016). Sorghum, a C4 plant, can also grow in adverse environmental conditions due in part to efficient water use and maintained photosynthetic activity under drought or high salinity conditions (Ratnavathi et al., 2011; Dar et al., 2018; Choudhary et al., 2021). This inherent resilience to a wide range of environmental conditions and differing stress tolerance mechanisms, and the ever-increasing need for developing cultivars capable of flourishing on resource-limited land unideal for more widespread crops, makes them prime candidates for comparative studies on environmental stress.

The abiotic factors selected for this study include drought, salinity and cold stress. The relevance and impact of these stressors on biofuel and agricultural crops has been well-characterized (Cramer et al., 2011; Koyro et al., 2012; Zhu, 2016; Zhang et al., 2022). Furthermore, there is evidence that the degree of salinity exposure can induce differential plant stress responses (Gupta and Huang, 2014). Thus, we employed two types of salt treatments for our experiments: a gradual increase in concentration labeled ‘salt stress’, and an immediate exposure to high concentrations termed ‘salt shock’. Chitosan, a biopolymer obtained by deacetylating chitin, a structural component of arthropods exoskeletons or fungal cell walls (Kean and Thanou, 2011), was also used as a stress treatment. While chitosan nominally falls within the biotic stress category due to triggering an immune response within plants (Chandra et al., 2015; Iriti and Varoni, 2015), application of this polysaccharide has in some cases been shown to enhance defense mechanisms and improve stress tolerance (Walker-Simmons et al., 1983; Hadwiger, 2013; Katiyar et al., 2015). On the other hand, it has also been suggested that chitosan can have a debilitating impact on root growth or microbe interactions, especially at higher concentrations (Calenn et al., 2018; Stasinka-Jakubas and Hawrylak-Nowak, 2022). We therefore applied chitosan to plants at a relatively high concentration (5 mg/ml) to observe the resulting effects. These abiotic and potential biotic stressors were selected to better understand how sorghum and camelina respond and adapt to different environmental conditions.

In this study, we employ a multi-modal approach to capture a wide range of physiological and biochemical responses of camelina and sorghum in response to environmental stress. Measurements of photosynthetic efficiency and leaf health parameters were captured using the MultispeQ device (Kuhlgert et al., 2016), while gas exchange and water-use efficiency was monitored with a LI-6800 portable photosynthesis system (Long and Bernacchi, 2003). Tissue electrical impedance, an indicator of water status and transport efficiency (Flores et al., 2019), was quantified with and on-plant biosensor (Multi-PIP, Growvera Inc.). Finally, biochemical shifts in plant tissues that could be indicative of stress responses, such as changes in cell wall composition or secondary metabolites (Lu and Rasco, 2010) were captured with Fourier transform infrared (FTIR) reflectance spectroscopy (HYPERION 2000, Bruker Inc.). These tools were chosen for their capacity to non-invasively monitor key physiological and biochemical markers of stress in living plants in real-time, facilitating the identification of stress signatures with high temporal resolution. Together, this multi-modal approach provides a multi-layered dataset that comprehensively combines multiple aspects of plant responses to environmental stress, offering avenues by which to detect unique and early signs of stress. This data in turn may inform future breeding programs and agronomic strategies, improving crop tolerance in a rapidly changing climate.

## Materials and Methods

### 2.1 Plant Material and Growing Conditions

*Sorghum bicolor* (Silage Master) and *Camelina sativa* (spring variety) seeds were obtained from the New Mexico State University Agricultural Science Center (Los Lunas, New Mexico, USA). Seeds were sown in 2.5-inch pots containing Metro-Mix 820 potting soil (Sun Gro Horticulture, Seba Beach, AB, Canada) and grown under full-spectrum AC120V LED grow lights (Honeywell International, Inc., Charlotte, NC, USA) on a 16-8-hour light/dark cycle with a commercially available timer. Plants were watered regularly with deionized (DI) water and kept at a constant temperature of ∼24°C with controlled humidity.

After four weeks of growth, sorghum had between 3 and 4 leaves, and camelina had between 8 and 9 leaves. Plants were divided into six treatment groups with four plants in each group. The experiment was repeated, and results from both were combined for data analysis.

Grape plants were grown at the Milagro Vineyards in Rio Rancho, New Mexico. European hazelnut (*Corylus avellana*) trees (cv. ‘Jefferson’) were grown in potted soil in an open-air greenhouse.

### 2.2 Treatment Conditions

The experimental treatments included the following: Control: plants watered with 500 mL of standard DI water. Chitosan: plants received 500 mL of a 2 mg/mL solution of chitosan (ThermoFisher Scientific). Drought: plants were watered with 500 mL of 10% polyethylene glycol (PEG) 8000 solution (ThermoFisher Scientific). Salt stress: plants were initially treated with 50 mM NaCl, and the salt concentration was gradually increased to 150 mM in increments of 50 mM every 12 hours. Salt shock: plants were immediately exposed to 150 mM NaCl. Cold: plants were moved from ∼24°C to a 4°C cold room and watered with 4°C DI water.

The total experimental duration was 14 days, with two days of gathering baseline physiological data before plants were exposed to environmental stress followed by 12 days of data gathering post treatment. Treatment solutions were replaced every 2 days.

### 2.3 Phenotypic Measurements

Photographs of the plants were taken at a fixed distance using a Logitech C930E USB camera. ImageJ software (Schindelin et al., 2012) was used to quantify changes in leaf area, leaf length, shoot length over time. Leaf pigmentation was calculated using the RGB values from the Histogram function. At the end of the experiment, plants were removed from soil and both the weight and length of the roots and shoots were measured. Plants were then individually wrapped in foil and dried in an oven at 80° C for ∼48 hours. Plants were removed from oven and the roots and shoots were reweighed.

Relative Leaf Water Content was quantified using the formula:

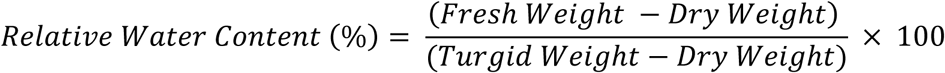

as described by Pieczynski et al. (2013).

### 2.4 Gas Exchange Measurements

Changes to gas exchange within plant leaves was quantified using two analytical tools: MultispeQ (PhotoSynQ Inc., East Lansing, MI, USA) v 2.0 and the LI-6800 Portable Photosynthesis System (LI-COR Biosciences, Lincoln, NE, USA) equipped with a 6800-01A leaf chamber fluorometers and 6 cm^2^ apertures.

The methodology for using the MultispeQ has been previously described (Kuhlgert et al, 2016). Briefly, as the sorghum leaves were narrower than the device sensor, following instructions listed on the PhotosynQ website, the MultispeQ was outfitted with a “mask” (a small piece of black construction paper) to restrict the optical window. After activating the MultispeQ through the PhotoSynQ app, the device was gently clamped onto a leaf, holding the device such that it remained upright and did not twist the leaf. A panel of environmental conditions and photosynthetic parameters were automatically measured over an ∼20 second period by the Photosynthesis RIDES 2.0 open-close protocol. Measured parameters include Phi2: the ratio of PSII quantum yield (*Φ*_PSII_) over ratio of incoming light; linear electron flow (LEF); and non-photochemical quenching (NPQt). After a measurement was completed, the device was removed from the leaf and the process was repeated for the next sample.

Methods for using and applying the LI-6800 have been previously described (Riches et al., 2020). Briefly, system warmup tests were run before each day’s experiments per the manufacturer’s recommendations. After the warmup tests were completed, the infrared gas analyzers (IRGAs) were matched using the range match feature. Survey measurements were taken by clamping the LI-6800 leaf chamber onto a leaf, waiting for the leaf to acclimate to the leaf chamber conditions, collecting gas measurements, and then removing the sample from the chamber and moving on to another leaf. Measurements were made at constant chamber air flow (600 µmol s^-1^), “auto” pump settings, chamber fan mixing speed (10,000 rpm), chamber air temperature (T_air_) (27 °C, save for the cold-treated plants which were kept at 4°C), constant relative humidity (60%), parabolic aluminized reflector (PAR) level (1500 µmol m^-2^ s^-1^), and carbon dioxide concentration (420 ppm).

Leaves measured were the two youngest, fully expanded, intact leaves for each plant, the 2^nd^ and 3^rd^ leaf for each sorghum plant and the 4^th^ and 5^th^ leaf for each camelina plant. Both tools were used daily between 9 a.m. and 1 p.m., measuring two fully expanded leaves per plant across three plants per treatment group.

### 2.5 Impedance Measurements

Impedance was measured every 15 minutes at frequencies of 1 or 100 kHz using the High Modes of two Multi-PIP (Growvera Inc., Albuquerque NM) impedance analyzers for sorghum and camelina. Each Multi-PIP impedance analyzer was attached to plants using two 3 m SMA-SMA cables connected to Growvera plant impedance probes (PIPs). These probes consisted of fine gauge, 200 mm long SMA to UMCC cables that connected to spring loaded clamps containing two 6-needle metal arrays with ∼1 mm long needles that are spaced ∼1 cm apart with no clamp material covering the plant tissue between the probes. Impedance is measured between the two needle arrays in each probe. For measuring impedance, clamps were gently clipped onto leaves (3^rd^ for sorghum, 6^th^ for camelina) such that each metal prong was inserted into and/or through the tissue. For camelina, the clamp was attached at the bottom right edge of the leaf from the stem. For sorghum, due to the size of the leaves, microneedles were inserted through the center of the leaf for maximum stability, although care was taken not to puncture the midrib. Once attached, the SMA cables were taped down to prevent unwanted twisting and minimize the amount of weight being placed on the leaves from the clamps. Once affixed, the Multi-PIP devices were connected to a power source and left to run autonomously. Multi-PIP data-collecting was reset every day to check for any potential issues. One sorghum and one camelina plant from each treatment group was measured using the Multi-PIP devices for each experimental run, as a single Multi-PIP can connect to a maximum of 8 probes at a time. Plants attached to the Multi-PIP did not undergo fluorescence, gas exchange, or FTIR measurements to avoid jostling the clips.

For grapevines, probes were attached to the stems or leaves and impedance was measured every 30 minutes at a frequency of 1 kHZ. For hazelnut trees, impedance sensor was installed by clamping it around the stem and gently squeezing the clamp to ensure the prongs fully penetrated into the stem. Impedance was measured every 30 minutes at a frequency of .1 kHz.

### 2.6 Psychrometer measurements

To investigate the relationship between impedance and stem water potential (Ψ), an impedance sensor Multi-PIP and a stem psychrometer (PSY-1, ICT-International, Australia) were both installed on a potted European hazelnut tree. The sensors were installed about 10 cm apart ∼30 cm above the potting soil and in the main stem. For the psychrometer, the installation procedure was conducted using the materials and instructions provided by ICT international. In short, the installation site was prepared using a razor blade to shave away the bark, cambium, and phloem which left a flat surface of sapwood/xylem larger than the chamber well exposed. The site was cleaned with DI water and then the psychrometer was clamped into place. The remaining exposed surface was covered with grease and then the psychrometer was insulated using polyester foam which was then covered by aluminum foil.

Data was collected every 30 minutes for a period of around 4 weeks. During the 4 weeks, trees started without water stress, water was withheld until severe water stress was evident, and then the tree was re-watered. The experiment was repeated 3 consecutive times using the same psychrometer. Hazelnut trees were used to overcome the challenges of using stem psychrometers on monocots, whose living stem tissues can contaminate and eventually damage the PSY1 thermocouple, and herbaceous plants, whose lack of woody stems can also pose significant installation challenges and limit accurate data collection.

### 2.7 FTIR Spectroscopy

The spectroscopic signature of plant leaves was quantified with a Bruker HYPERION 2000 Optical Microscope (Bruker Corporation, Billerica, MA) using a Mercury cadmium telluride (MCT) detector with a 64 x 64 focal plane array (FPA) and the OPUS software Version 7.2.139.1294 (Bruker Corporation, Billerica, MA). As the thickness of living, intact plant leaves blocked the light necessary to generate transmission spectra, the reflectance spectra of whole, intact leaves were measured. Background FTIR reflectance spectra was measured using the silver mirror built into the optical microscope stage. Background was then automatically subtracted from sample measurements. Once this baseline has been established, the requisite amount of each sample was placed on the Hyperion microscope stage, whereupon a visual image was recorded using the Video Wizard program using a 4x objective. Each measurement was taken using a spectral range of 8000–600 cm^−1^ at a spectral resolution of 36 cm^−1^ and sixteen co-added scans, adding up to ∼6 minutes of measurement time per sample. Two leaves from each plant were measured each day, for a total of n=16. After the experiment had concluded, three leaves and three roots of each plant were measured using reflectance. After whole plants were dried, the reflectance measurements were repeated on the dried leaves. FTIR spectra were visualized using the open-source program Spectragryph Version 1.2.16.1 (Menges, 2024). Data processing was performed by converting reflectance to 1/log(reflectance), baseline corrected using Spectragryph’s “Advanced Baseline” adaptive method with 15 coarseness and 0 offset, and then smoothed using a Savitzsky-Golay filter with an interval of 10 and a polynomial order of 3. Data was then normalized by dividing values by the maximum value within the spectra.

### 2.8 Statistical Analysis

One-way ANOVA followed by Tukey’s Honestly Significant Difference (HSD) post hoc tests were used to determine significant differences (p = 0.05) among the treatments. Normality and homogeneity of variances were checked using the Kolmogorov-Smirnov and Levene’s tests, respectively. Significant outliers were identified using the Mahalabinois distance and removed. Principal Component Analysis was performed in Python using the “sklearn” package (Pedregosa, 2011). All code was developed in Python 3.11.7 Version 1.4.02 of the instrument software for the work reported here. Development of code was aided using ChatGPT. Code is publicly available through GitHub. Impedance data was graphed and analyzed using the “Multi-PIP Analysis Suite”, an open-source Python module developed and provided by Growvera.

## Results

### 3.1 Growth and Morphological Responses to Stress

Twenty-four camelina and twenty-four sorghum plants were organized into six groups with four camelina and four sorghum seedlings each. After gathering baseline data for two days, each group was exposed to either control, chitosan, cold, drought, salt stress, or salt shock conditions for a period of twelve days, with physiological data gathered throughout. The experiment was repeated twice. At the end of the treatment period, visual assessment indicated drought and salinity triggered drastic morphological changes within camelina seedlings, while cold and chitosan seemingly reduced or increased shoot length, respectively (Supplementary Fig. S1). For sorghum, (Supplementary Fig. S2), cold triggered severe morphological changes, drought and salinity conditions had a less severe but still noticeable effect, and chitosan may have increased leaf length. When growth indices were quantified, data regarding camelina seedling growth traits (Table 1) indicated significant differences between un-stressed and stressed plants. Shoot length (SL) and fresh shoot mass (FSM) were significantly reduced in response to drought, salinity, or cold. Dry shoot mass (DSM) was reduced in response to drought and salt stress. Leaf water content (LWC) decreased in response to both drought and salinity. In contrast, chitosan treatment triggered an increase in both SL and fresh root mass (FRM).

**Table 1:** Growth indices of camelina seedings exposed to environmental stress.

| Camelina | Length (mm) |  | Fresh Mass (mg) |  | Dry Mass (mg) |  |  | Leaf Water Content (%) |
| --- | --- | --- | --- | --- | --- | --- | --- | --- |
|  | Shoot | Root | Shoot | Root | Shoot | Root | Root:Shoot |  |
| Control | 233±29 | 172±31 | 861±209 | 30±13 | 109±24 | 22±10 | 0.20±0.07 | 86±2 |
| Chitosan | 283±35* | 188±49 | 1102±253 | 53±14** | 133±34 | 31±12 | 0.24±0.07 | 85±1 |
| Cold | 175±25** | 201±45 | 563±236* | 47±15 | 108±54 | 33±8.7 | 0.36±0.15 | 81±4 |
| Drought | 134±39** | 116±43 | 71±52** | 13±7.1 | 40±25** | 12±6.8 | 0.40±0.33 | 25±7** |
| Salt Stress | 163±17** | 140±41 | 129±20** | 20±12 | 51±30* | 17±12 | 0.36±0.22 | 41±9** |
| Salt Shock | 183±21* | 163±21 | 107±34** | 24±7.1 | 58±17 | 17±5.7 | 0.28±0.06 | 37±19** |

± =SD, \* = p<.05, \*\* = p<.01, n = 7 plants

The sorghum seedling growth indices (Table 2) exhibited less severe differences between stressed and un-stressed plants. FSM was only reduced in cold-treated plants, with SL or SDM not approaching the conventional limits of significant for any stress group. However, LWC did show significant differences, exhibiting high reductions in response to cold, drought, or either salinity treatment as compared to plants grown under normal environmental conditions.

**Table 2:** Growth indices of sorghum seedings exposed to environmental stress.

| Sorghum | Length (mm) |  | Fresh Mass (mg) |  | Dry Mass (mg) |  |  | Leaf Water Content (%) |
| --- | --- | --- | --- | --- | --- | --- | --- | --- |
|  | Shoot | Root | Shoot | Root | Shoot | Root | Root:Shoot |  |
| Control | 250±79 | 216±42 | 393±270 | 35±20 | 54±38 | 22±17 | 0.42±0.19 | 93±5 |
| Chitosan | 302±28 | 232±56 | 587±94 | 58±16 | 71±9.1 | 31±9.6 | 0.43±0.10 | 97±2 |
| Cold | 199±50 | 207±62 | 144±116* | 23±15 | 31±19 | 12±6.9 | 0.44±0.23 | 18±10** |
| Drought | 232±43 | 208±77 | 195±108 | 65±43 | 47±20 | 41±23 | 0.86±0.27** | 24±19** |
| Salt Stress | 234±36 | 175±42 | 228±74 | 49±6.1 | 36±14 | 17±8.2 | 0.48±0.12 | 38±27** |
| Salt Shock | 264±40 | 204±34 | 170±93 | 51±23 | 49±23 | 23±8.2 | 0.52±0.12 | 31±24** |

± =SD, \* = p<.05, \*\* = p<.01, n = 7 plants

When SL, leaf length (LL), and leaf color (LC) of the seedlings were measured over time (Fig. 1), the SL of camelina seedlings exposed to abiotic stress (Fig. 1A) began to deviate at ∼day 7, becoming significantly distinct by day 9. For LL (Fig. 1B), cold and chitosan exposure did not cause significant changes, while the drought-treated plants became significantly reduced by day 3 and the salt-treated plants by day 4. For LC (Fig. 1C), the stressed plants had a slight decrease in overall pigmentation but was not significant. However, when the green color intensity (GC) was quantified (Supplementary Fig. S3A), drought and salt significantly reduced green pigmentation by day 6 and cold by day 10. For sorghum, measured growth indices once again showed little deviation between treatments over time. SL (Fig. 1D) and LL (Fig. 1E) never reached significance thresholds. LC (Fig. 1F) did significantly decrease in abiotically-stressed plants by day 10, however, this reduction was not observed in GC (Supplementary Fig. S3B).

**Figure 1.**
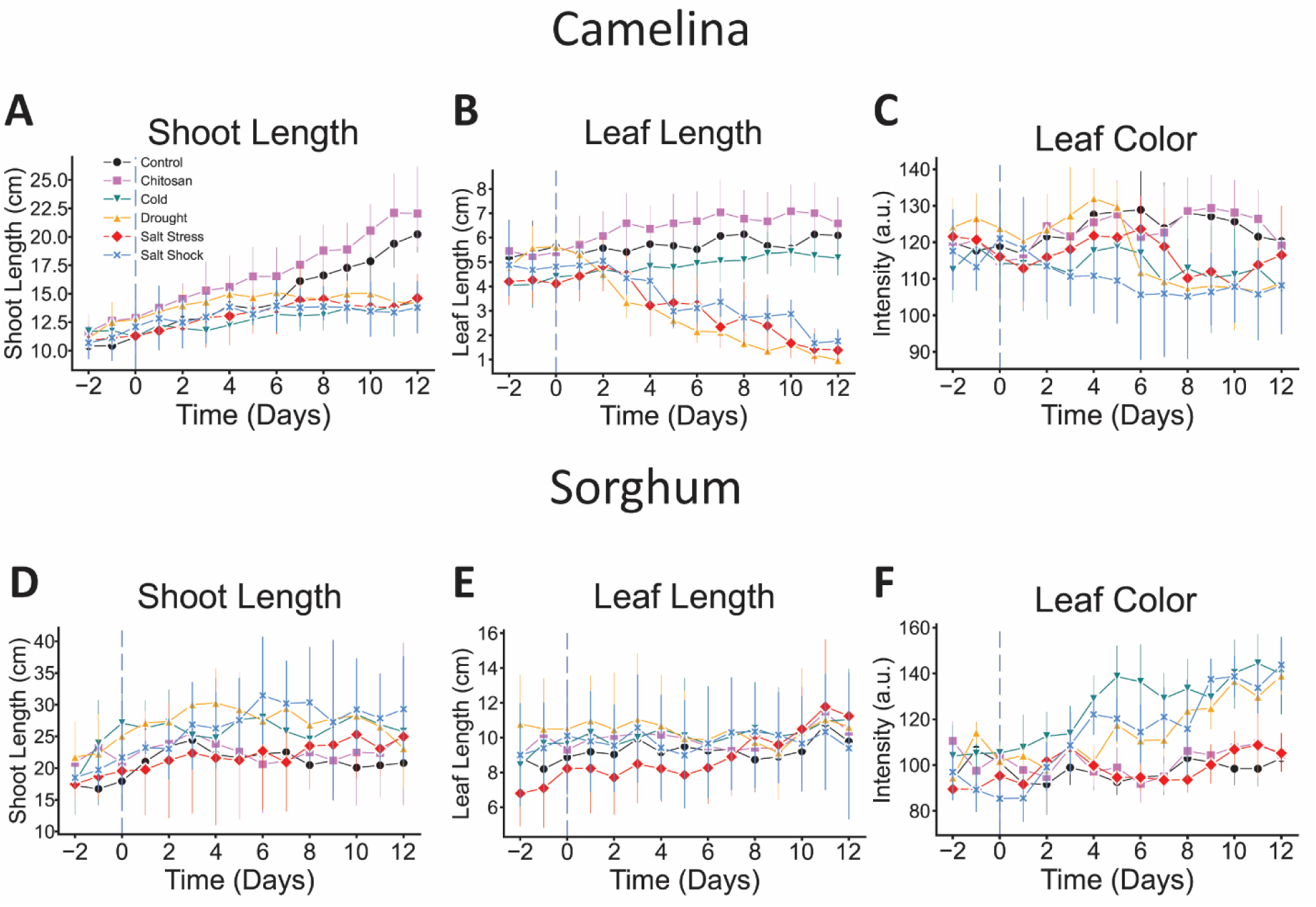
Growth indices of camelina and sorghum exposed to environmental stress over time. (A and D) Shoot length, (B and E) leaf length, and (C and F) leaf color intensity of camelina and sorghum seedlings exposed to control, chitosan, cold, drought, salt stress, or salt shock was recorded every day of a 14 day experimental period. Vertical blue dashed lines indicate when treatment began. Error bars indicate standard deviation. n=7 plants (shoot length) or 12 leaves (leaf length and leaf color).

### 3.2 Fluorometric and Gas Exchange Responses

Highly significant variations (P≤0.01) were recorded when the gas exchange characteristics of abiotically-stressed camelina and sorghum seedlings were compared to normally grown plants (Fig. 2). Overall, the parameters for chitosan and cold-treated plants remained stable and displayed little deviation from non-stressed plants. In contrast, drought and salt conditions caused major alterations to all measured variables. Drought imposition induced changes in non-photochemical quenching (NPQt) (Fig. 2A), chlorophyll fluorescence (ΦPSII) (Fig. 2B), linear electron flow (LEF) (Fig. 2C), photosynthetic rate (A) (Fig. 2D), transpiration (E) (Fig. 2E), and stomatal conductance (gs) (Fig. 2F), all becoming significantly different from control plants 2 or 3 days posttreatment. Salt stress and salt shock-treated plants exhibited similar phenotypes, although they only became significant 4 or 5 days posttreatment.

**Figure 2.**
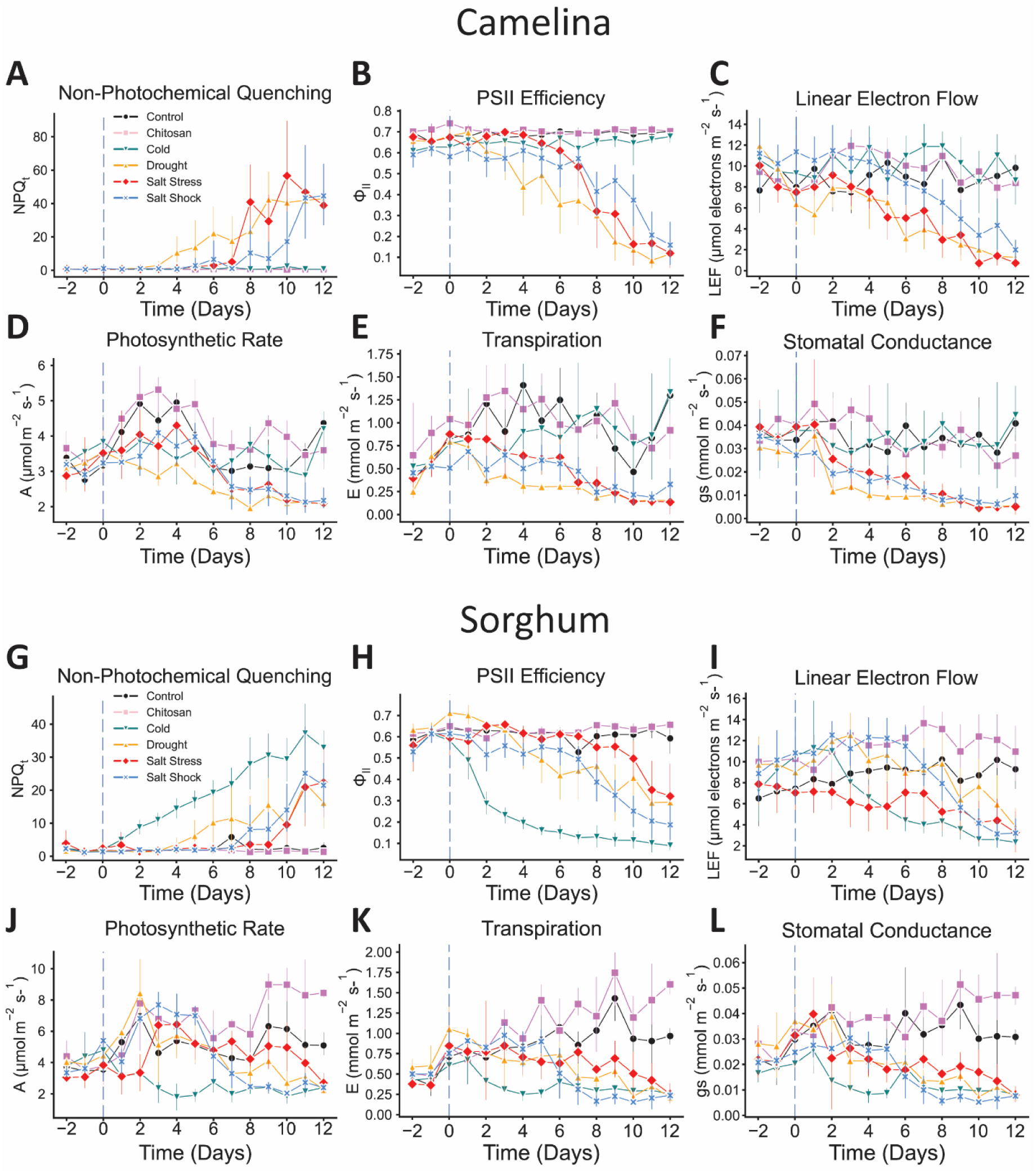
Fluorometric and gas exchange parameters of camelina and sorghum exposed to environmental stress over time. (A and G) Non-photochemical quenching, (B and H) PISII efficiency, (C and I) linear electron flow were measured in camelina and sorghum plants undergoing environmental stress using a MultispeQ. (D and J) Photosynthetic rate, (E and K), transpiration, and (F and L) stomatal conductance were measured using an LI-6800. Vertical blue dashed line indicates when treatment began. Error bars indicate standard deviation. n=12 leaves.

Chitosan-treated sorghum plants differed little from the control. However, plants subjected to cold stress displayed a rapid and significant decline in all measured gas exchange parameters, starting 1 to 2 days after the initiation of the cold treatment. Drought and salinity resulted in slower, more gradual deviations from the control. For these stress treatments, ΦPSII, LEF and A began to decline after several days of exposure, with the most pronounced effects seen after day 7, while non-photochemical quenching (NPQt) exhibited the opposite response. E and gs also showed gradual reductions under both drought and salt treatments, with salt shock-treated plants showing a slightly more rapid decline than salt stress-treated plants.

### 3.3 Impedance Measurements

Electrochemical impedance spectroscopy has recently emerged as a rapid, non-destructive method to directly quantify plant parameters related to health and stress in real-time (Zhang and Willison, 1993; Jócsák et al., 2019; Van Haeverbeke et al., 2023). Several groups have developed modalities for measuring plant impedance, but these tend to be limited by large measuring probes, restrictions to woody tissues in mature plants, or an inability to differentiate between internal structures (El Khaled et al., 2017; Ibba et al., 2020, Basak et al., 2020; Garlando et al., 2022; Reynolds et al., 2023). Recently, Reed et al. published a method for quantifying impedance in plants via wearable biosensors consisting of <1 mm microneedles, which were capable of quantifying and detecting unique impedance patterns in a variety of tissues (stem, lamina, petiole). The setup described by Reed et al. was then further developed into a small, portable, low-cost impedance analyzer for continuously monitoring plants in the field, known as the Multi-PIP (Supplementary Fig. S4). When impedance measurements obtained using the Multi-PIP device were compared with concurrent water potential measurements from a psychrometer during a drying trial (Supplementary Fig. S5), the data demonstrated a correlation between impedance and water potential fluctuations across days, supporting the ability of the Multi-PIP system to reflect changes in plant water status over time. In addition, when the Multi-PIP was used to quantify impedance in vineyard grapes over a growing season, key events, such as bud break or initiation of watering, were reflected in significant changes to impedance values (Supplementary Fig. S6). This data supports the hypothesis that impedance can reflect physiological changes in live plant tissue in response to alterations in environmental conditions. Based on these findings, we therefore sought to measure impedance within living, intact camelina and sorghum leaves in response to severe and sustained environmental stress (Fig. 3).

**Figure 3.**
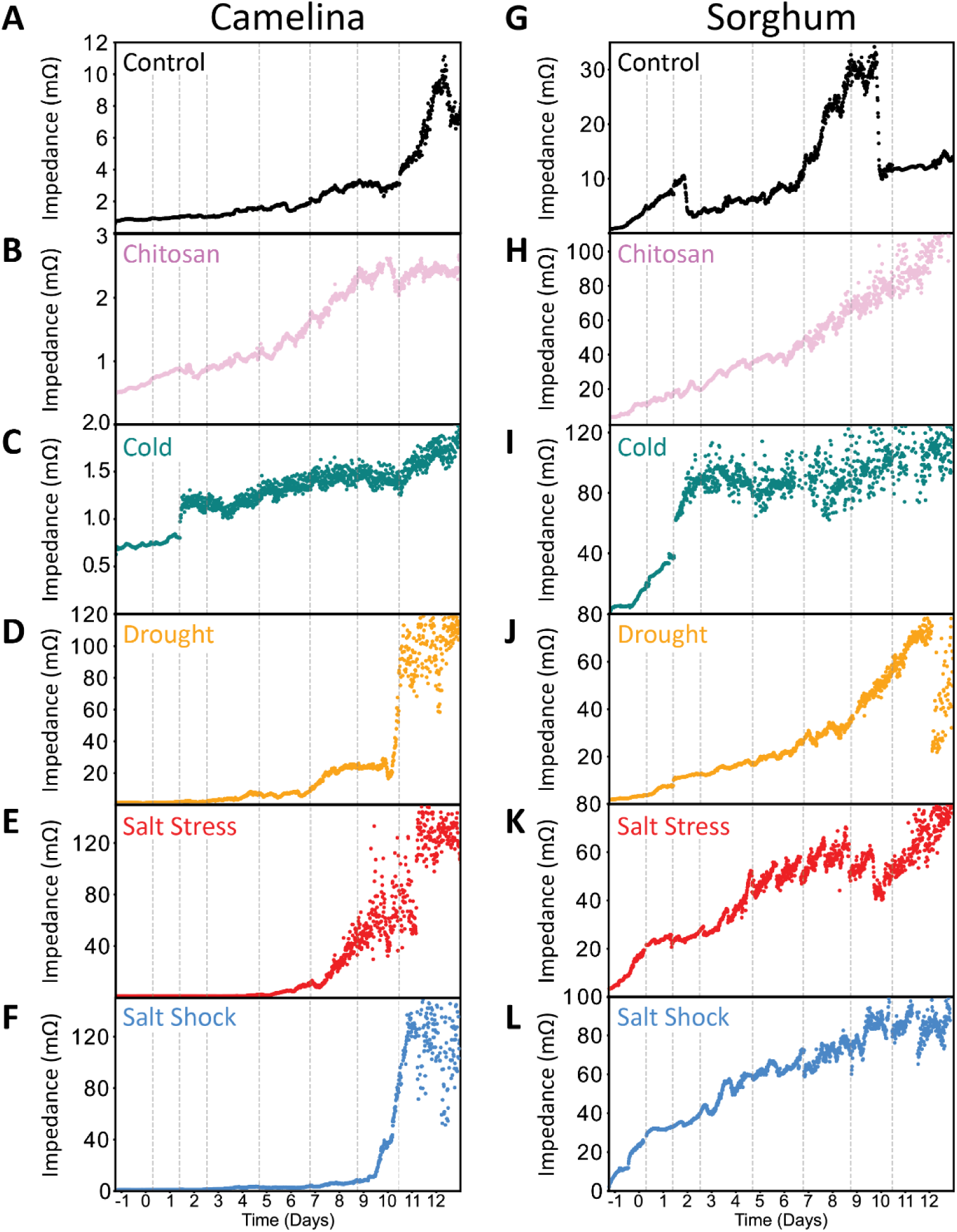
Impedance measurements of camelina and sorghum exposed to different types of environmental stress. Camelina and sorghum seedlings were exposed to either (A and G) control, (B and H) chitosan, (C and I) cold, (D and J) drought, (E and K) salt stress or (F and L) salt shock, and impedance of select leaves was measured using the Multi-PIP at 1000 Hz frequency. Y-axis is impedance measurements in mOhms. Vertical dashed lines indicate when treatment began (first line) and when the treatment solution was renewed (following lines).

#### 3.3.1 Camelina Impedance Responses

For control camelina (Fig. 3A), impedance measurements remained relatively low throughout the measurement period, rising rapidly only after day 10. However, as this spike was not observed in other control, chitosan, or cold-treated plants, it is likely an erroneous state caused by the clips losing contact with the leaf tissue, possibly due to plant growth or human error. Chitosan-treated plants (Fig. 3C) gradually increased in impedance over time, with readings below 3 mOhms for the entirety of the experiment. Cold-treated camelina (Fig. 3E) displayed a notable increase in variability after one day of treatment, as evidenced by the scatterplot shifting from a nearly linear configuration to a thicker band. However, impedance overall exhibited little change in value beyond known dips associated with diurnal patterns in water dynamics. In contrast, drought and salt-treated plants (Fig. 3G, 3I, 3K) displayed far greater fluctuations, as maximum impedance values were one order of magnitude greater than the previous three treatment groups. We observed the following: impedance of drought induced stress increased on day 4, day 7, and a large spike on 10; impedance gradually increased in response to salt stress starting on day 5; and impedance also increased due to salt shock induction on day 7 and spiked on day 9.

When the data was analyzed to emphasize early impedance response to environmental stress (Supplementary Fig. S7), drought induction resulted in a rapid impedance increase a few hours after initial treatment, while salt shock and salt stress triggered surges a few days after exposure. Similar patterns and trends were observed when the experiment was repeated at a higher frequency (100 kHz versus 1 kHz) (Supplementary Fig. 8).

#### 3.3.2 Sorghum Impedance Responses

Notably overall sorghum impedance values were higher in comparison to those measured in camelina, possibly due to differences in plant morphology. Control (Fig. 3B) values exhibited slightly higher levels of variability, but overall remained below 35 mOhms, while chitosan-treated (Fig. 3D) plants exhibited relatively a constant increase over time. Cold-treated sorghum (Fig. 3F) displayed a rapid escalation in impedance immediately after treatment began. Sorghum plants exposed to drought and salt conditions (Fig. 3H, 3J, 3L) exhibited more gradual increases in impedance over time. Drought stricken plants displayed a slow, steady rise in impedance starting around day 3, whereas salt stress and salt shock treatments caused distinct surges beginning after day 6. Notably, the diurnal changes in impedance were retained within the salt stressed and salt shocked plants when compared to those undergoing drought.

However, unlike camelina wherein impedance values remained relatively consistent for the two days prior to treatment, sorghum plants exhibited sharp impedance increases before environmental conditions were altered (Supplementary Fig. 7), obscuring potential early impedance changes in response to abiotic or biotic stress within these seedlings. Regardless, when the experiments were repeated with a frequency at 100 kHz (Supplemental Fig. 8), the same general trends in response to each type of stress were observed, once again suggesting that the impedance of plant leaves is altered in response to environmental stress.

### 3.4 FTIR Spectral Analysis and Principal Component Analysis (PCA)

To investigate potential biochemical changes occurring in camelina and sorghum in response to stress treatments, Fourier-transform infrared (FTIR) spectroscopy was conducted. Previous application of FTIR to plant leaves mainly utilized dried and homogenized tissue (Naumann et al., 2010; Lu and Rasco, 2010; Cen and He, 2007), limiting the scope of information that could be obtained. Reflectance measurements of non-treated intact leaves of both camelina and sorghum (Supplementary Fig. S9) captured 15 features with highly similar peak location and amplitude between species. These peaks were assigned to specific functional groups and compounds commonly found in plant tissues (Supplementary Table 3) based on (Nikalje et al., 2019; Ertani et al., 2018; Topală and Rusea, 2018; Yang and Yen, 2002). Some key assignments include the 3519/3488 peak (OH/NH stretching), associated with alcohols and various polysaccharides (e.g., cellulose), 1753/1744 cm⁻¹ (C=O stretching) found in ester groups and linked to compounds such as pectin, lignin, suberin, and cutin, and 820/816 cm⁻¹ (out of plane CH bending), representing aromatic rings, these peaks indicate the presence of crystallized cellulose within the sample.

After establishing baseline spectra, FTIR spectroscopy was conducted on leaf samples at the end of the 12-day experimental period (Fig. 4). Fig. 4A shows the FTIR spectra for camelina plants across all treatments. Camelina exhibited substantial spectral shifts in response to drought and salt, particularly in the protein and carbohydrate regions. Cold treatment had less of an impact, consistent with greater tolerance to cold stress, but there were still visible changes in the reflectance spectra of cold-stressed plants, most notably in the 3500 cm⁻¹ and 1200 cm⁻¹ regions. For sorghum (Fig. 4B), the most significant changes in the spectra were observed under cold and drought treatments, with notable alterations in the alcohols (3500 cm⁻¹) and protein (1700-1600 cm⁻¹) regions. In contrast, while both salt shock and salt stress exposed induced similar biochemical changes, they were less prominent overall, and plants treated with salt stress displayed overall greater spectral heterogeneity, with some more closely resembling the molecular spectra of control plants. When Principal Component Analysis was performed (Fig. 4C and D), the PCA plots revealed clear separations between the different treatments. For camelina (Fig. 4C), drought and salt led to the most significant divergence from the control, while cold-treated plants displayed minor but significant separation. For sorghum (Fig. 4D), drought and cold treatments resulted in the greatest divergence from the control, with distinct clustering observed along PC1. In contrast, salt shock displayed more intermediate clustering, while some of the salt stress spectra clustered with control, again displaying a wide range of heterogeneity within the biochemical profile. When examining which wavenumbers contribute the most to the variance observed in the PCA (Fig. 4E and F), for both camelina and sorghum, the alcohols (3500 cm⁻¹) and ring (800 cm⁻¹) regions were the primary drivers of the variance, while protein (1700-1600 cm⁻¹) and carbohydrates (1200-1000 cm⁻¹) played a lesser, but still significant role.

**Figure 4.**
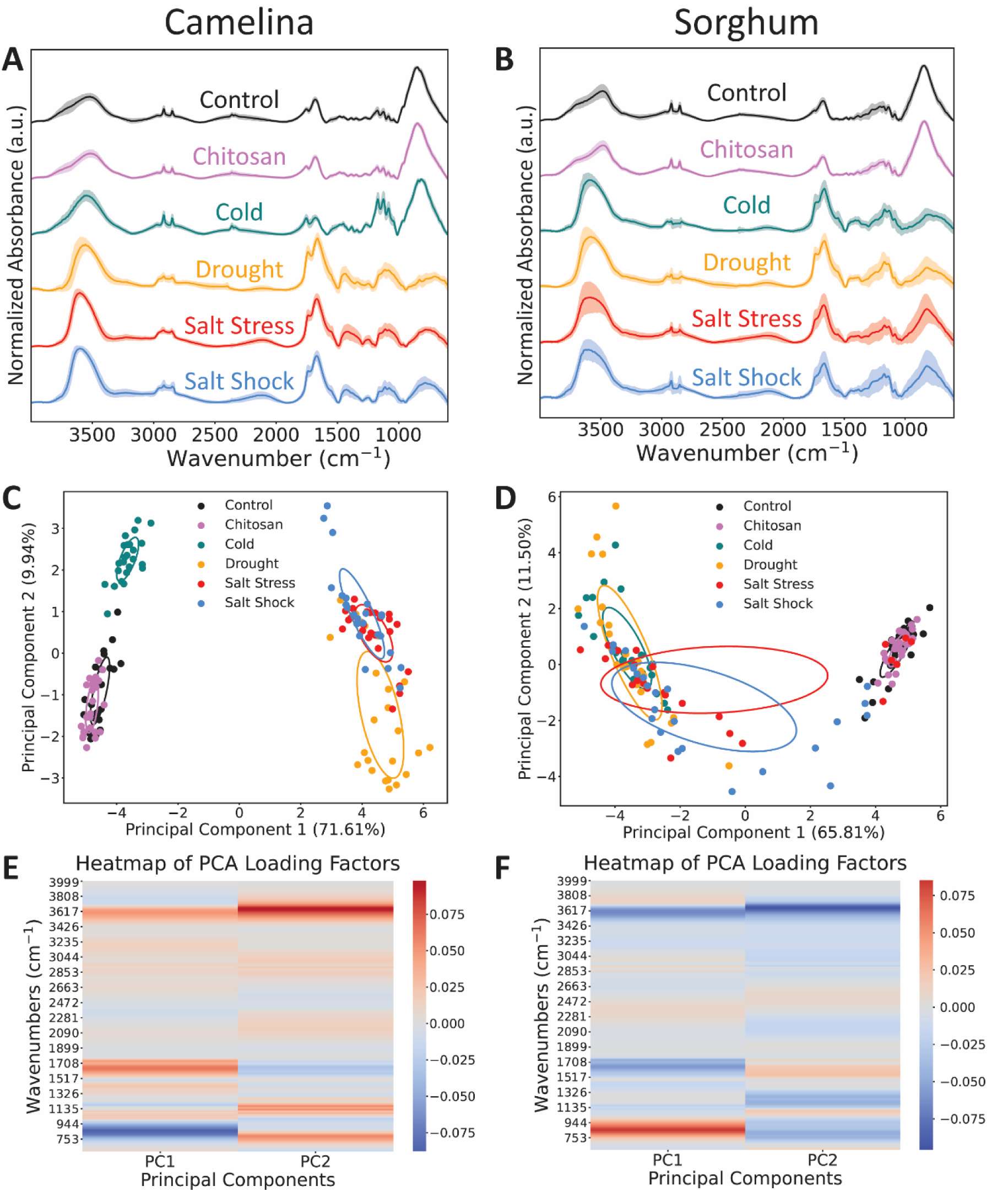
FTIR spectra of fresh camelina and sorghum leaves exposed to environmental stress. The reflectance spectra or stress or non-stressed (A) camelina and (B) sorghum plants were measured at the end of the experimental period. (C and D) Principal Component Analysis (C and D) and (E and F) heatmaps of PCA Loading Factors were also performed. n=24.

After measurements of the fresh leaves were concluded, samples were dried and FTIR reflectance spectra were again taken (Supplementary Fig S10). Notably, this sample preparation method resulted in the spectra of the control plants becoming indistinct from those of fresh plants, suggesting that the drying process may influence the FTIR spectra. The reflectance FTIR of fresh roots was also performed (Supplementary Fig. S11). While drought-treated roots displayed a unique spectra profile in comparison to unstressed roots, the drought-specific peaks were similar to those found in the reflectance spectra of pure PEG, suggesting these features may be due to PEG lingering on the surface of samples. Overall, these results suggest that not only are intact, unperturbed plants a viable target for spectroscopic detection of plant stress, but they may also in some ways be favorable compared to dried samples or other types of plant tissues as an indicator of stress response.

#### 3.4.1 Changes in FTIR Spectra in Response to Stress Over Time

After establishing that FTIR spectroscopy could detect spectral shifts in response to abiotic stress, the next step was to quantify the spatial dynamics of these changes. As such, FTIR measurements were conducted on leaves for every day of the 12 day experiment period. For camelina (Supplementary Fig. 12), the control and chitosan-treated camelina plants exhibited stable FTIR spectra throughout the experimental period, while the spectra of abiotically-stressed plants exhibited changes over time. The same could be observed for sorghum (Supplementary Fig. 13). When the rate of change of absorbance at each wavenumber were calculated, for camelina (Supplementary Fig. 14), control and chitosan exposure induced very little rate of change over time. In contrast, abiotically stressed plants exhibited much greater alterations, with absorption increasing quickly in the 3500 cm^-1^ region over time, in contrast to control and chitosan, where that region showed a small dip. In the lipid region (3000-2800 cm^-1^), abiotic induction absorption increased at a slower rate over the course of the experiment. Absorption in regions associated with proteins (1700-1600 cm^-1^) quickly increased for drought and salt stress, moderately increased for salt shock, and slightly decreased from cold exposure when compared to control plants. For regions associated with carbohydrates (1200-1000 cm^-1^), major increases were observed for cold, drought, and salt stress, and a moderate increase for salt shock. The absorbance at wavenumbers associated with crystallized cellulose (800 cm^-1^) decreased quickly over the course of the experiment for the abiotically stressed plants in comparison to minor decreases for control or chitin-exposed plants. Sorghum (Supplementary Fig. 15) exhibited much of the same changes, although notably cold-treated plants exhibited relatively minor shifts in comparison to the control or chitosan-treated plants, mainly in the alcohol-associated spectral region. Additionally, salt-stressed plants also showed a minor increase in this region, despite distinct changes in in other areas that resemble drought and salt shock in amplitude.

Finally, heat maps of the reflectance intensity across the course of the experiment shows precisely which days shifts began to occur. For camelina plants (Supplementary Fig. 16), cold treatment began to show significant changes by day 8 in regions associated with alcohols and carbohydrates. Drought showed changes in regions associated with alcohols on day 6 and significant changes in regions associated with alcohols, proteins, and cell wall components by day 8. Salt stressed plants indicated significant changes in these regions by day 6, while salt shock did not show shifts until day 11. In sorghum (Supplementary Fig.17), we observed that cold treatments induced changes by day 12. Drought began showed changes in regions associated with alcohols on day 6 and significant changes in regions associated with alcohols, proteins, and cell wall components on day 8. Salt stress induced changes in regions associated with alcohols as early as day 2 and significant changes in regions associated with alcohols, proteins, and cell wall components on day 11. Salt shock induced changes in regions associated with alcohols on day 3 and significant changes in regions associated with alcohols, proteins, and cell wall components on day 8. These results demonstrate that rates of change in FTIR spectral profiles over time in response to environmental stress can be used as distinct indicators of certain stresses imposed on the plants.

## Discussion

This study assessed the physiological and biochemical responses of four week old *Camelina sativa* and *Sorghum bicolor* plants to chitosan, cold, drought, salt stress, and salt shock exposure using a range of non-invasive measurement tools. Our findings reveal distinct stress response mechanisms between the two species, highlighting camelina’s pronounced sensitivity to drought and salinity and sorghum’s susceptibility to cold stress. These differential stress responses could be quantified as unique physiological profiles by utilizing a multi-modal approach.

Camelina was highly sensitive to drought and both types of salinity treatment. Under these conditions, reductions in growth indices and photosynthetic activity were rapid, usually taking place only a few days after abiotic stress exposure. FTIR and PCA analyses revealed significant biochemical changes, especially in the water, protein, and glycosidic bond regions, suggesting that camelina’s cellular integrity and water transport mechanisms are severely impacted in response to these types of environmental stress, a finding supported by previous literature (Gall et al. 2015; Zhu 2016; Baranza et al., 2021). In contrast, cold had little impact on gas exchange measurements but did deviate from control plants in growth indices, impedance readings, and FTIR spectra, suggesting that while major functions of these plants are not interrupted, the mechanisms required to adapt to cold environments consume resources that would otherwise be allocated towards growth.

In sorghum, cold stress induced significant reductions in key growth metrics such as leaf area and photosynthetic efficiency. This was coupled with pronounced changes in impedance, once again suggesting that environmental stress impairs water transport efficiency, leading to the degradation of cell wall components in plants. In contrast, responses to drought and salinity were more gradual, with observable changes in gas exchange and FTIR spectra only emerging later in the experiment. The changes observed in the lipid and protein regions of the FTIR spectra further suggest that sorghum plants undergo metabolic shifts as a means by which to increase abiotic stress tolerance, highlighting a potential target for detecting early signs of plant stress and developing more resistant crops overall.

Of all the treatments, application of chitosan seemingly had a positive effect. Previous research suggested that application of chitosan stimulates root growth to aid in nutrient and water absorption, and boosts chlorophyll content (Katiyar et al., 2015), which is in agreement with the enhanced root and shoot biomass, and photosynthetic activity seen in our experiments. In contrast, negative effects of chitosan had previously been documented in Arabidopsis. It is possible that higher concentrations of chitosan are required to produce deleterious effects in camelina or sorghum, though our study had higher chitosan concentrations than what is usually used (5 mg/ml). Regardless, this work reaffirms that chitosan is a beneficial supplement for bolstering overall plant growth and resistance to stress. Our findings on the differences between salt stress and salt shock exposure suggested that plants seemed to respond more rapidly to and have a worse outcome from gradual salt exposure rather than acute salt exposure. More research is needed, but this emphasizes a promising avenue regarding how minute changes to the magnitude or application of environmental stress can translate into detectable characteristics in the plant stress response.

One of the central objectives of this study was to evaluate the efficacy of rapid, non-invasive, and non-destructive tools for monitoring plant stress responses. While growth indices can be used to quantify changes to plant health in response to environmental stress over time, these measurements are laborious, time-intensive, and generally rely upon the plant exhibiting visual indicators of stress, at which point remedies are much more difficult. The use of technologies such as Multi-PIP impedance sensors, MultispeQ, and FTIR spectroscopy provided real-time insights into plant physiological changes, allowing for the early detection of plant stress in response to environmental conditions. The sharp declines in net CO₂ assimilation, stomatal conductance, and chlorophyll fluorescence under drought and salinity in camelina, and cold stress in sorghum, highlight the utility of gas exchange parameters in detecting stress-induced disruptions in photosynthesis. The correlation between these measurements and the FTIR spectra also provides a comprehensive view of how stress affects both plant physiology and biochemistry. Multi-PIP impedance sensors proved to be highly effective in capturing real-time changes in tissue water content and cellular integrity without disturbing the plant. The rapid increase in impedance (on the order of hours) observed under drought and salinity in camelina, suggests that impedance measurements could be a reliable indicator of early stress response, particularly under extreme environmental conditions. This is also supported by the clear deviations between environmental treatments in the FTIR spectra regions associated with OH bending, further demonstrating that water can be linked to specific stressors. FTIR spectroscopy also allowed for the non-invasive analysis of a wide variety of biochemical changes. The spectral shifts observed under salt shock, drought, and cold treatments revealed alterations in carbohydrates, lipids, and proteins, which are known to play critical roles in plant stress adaptation. Importantly, several of these regions displayed unique alterations on a temporal or amplitude scale depending on the type of stress induction, highlighting the ability of IR spectroscopy to probe plant stress response in real-time. Furthermore, many of these treatment-induced changes disappeared or became difficult to distinguish in either dried or non-leaf samples, suggesting that this tool may be ideal for assessing living, easily accessible plant tissues for signs of plant stress.

While this study provides valuable insights into plant stress responses and the applicability of both established and novel tools to capture these dynamics, there are limitations to the approaches employed. The use of polyethylene glycol (PEG) to simulate drought conditions may have introduced artifacts that do not perfectly replicate field drought scenarios. Future studies should consider using water deprivation methods that better reflect natural drought conditions. It should also be noted that plants undergoing Multi-PIP measurements were excluded from gas exchange and fluorescence measurements to avoid detaching the probes, and in turn it was found these plants displayed notable increases in growth indices (see Supplementary Datasheet 1).

Considering that repeated clamping of gas exchange chambers has been found to cause damage to leaf tissue (Marler and Mickelbart, 1992), it is possible that the frequent handling of plants for repeated physiological measurements in this study may have also introduced latent forms of stress. Future experiments should seek less invasive measurement devices, such as those utilizing imaging or that don’t need repeated contact. However, it should also be noted that one of the tools that does not require contact, the Multi-PIP, occasionally exhibited unexpected increases in impedance during the experiment, most notably within sorghum leaves. We attribute these spikes to probe-tissue contact becoming loose over time, likely due to growth or the initial small size of leaves, as impedance measurements in stem tissue of hazelnut or grapevines did not exhibit these patterns. As such, more rigorous methods of attachment or more stable tissues would likely resolve this issue. Moving on, most of these tools used in this study were unable to provide spatial information with a high quality of resolution, greatly hampering our insight into how plant physiological processes are altered in response to stress at the tissue, cellular, or subcellular level. This aspect of data gathering is essential to fully understanding the mechanisms by which plants deal with environmental stress and may provide both key insights into how early signs of stress response deviate and potential offer molecular and morphological targets for gene-editing to develop plants more resistant to environmental stress. A final limitation is that the controlled environment of the study does not fully represent the complex interactions that occur in field conditions, such as fluctuating environmental factors or pest pressures. Scaling these non-invasive tools for large-scale field applications will also require improvements in sensor robustness, data integration, and deployment across larger agricultural landscapes.

## Conclusion

In this study, we compare multiple non-invasive tools for early detection of common plant stresses and demonstrate the value of multi-modal approaches to highlight the divergent strategies employed by sorghum and camelina to cope with abiotic stress, including potential avenues for detecting early plant stress response to environmental conditions or correlating specific physiological changes to unique environmental parameters. As climate change continues to challenge global food security, adopting smart technologies for crop monitoring will be critical for enhancing resilience and maintaining productivity in a sustainable manner.

## Acknowledgments

This material is based upon work supported by the U.S. Department of Energy, Office of Science, Biological and Environmental Research (BER) Biomolecular Characterization and Imaging Science Program (BCIS) under Award number [LANLF31W]. This work was performed, in part, at the Center for Integrated Nanotechnologies, an Office of Science User Facility operated for the operated for the U.S. Department of Energy (DOE) Office of Science by Los Alamos National Laboratory (Contract 89233218CNA000001) and Sandia National Laboratories (Contract DE-NA-0003525). LA-UR-24-31667.

## Author Contributions

All authors participated in experimental design. K.S., S.B, and E.P collected data. K.S., S.B., and E.P created figures. K.S. and D.M. developed code and wrote manuscript. D.M, D.R., R.N., D.H., and J.W. provided editing and feedback.

