## Supplemental Figures 1-17 and Supplementle Table 1 for "Characterizing the Responses of Camelina and Sorghum to Environmental Stress through a Multi-Modal Approach"

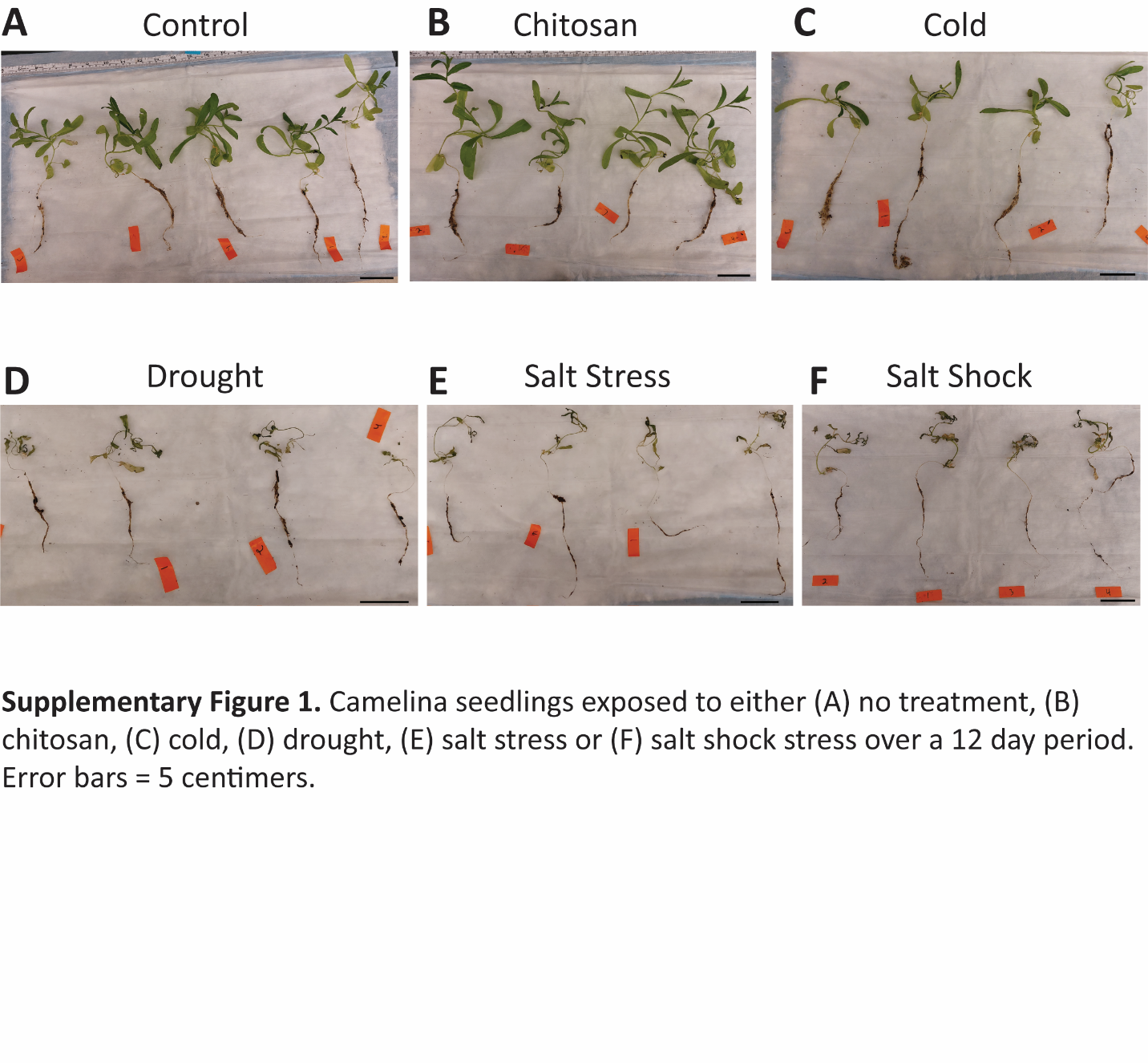

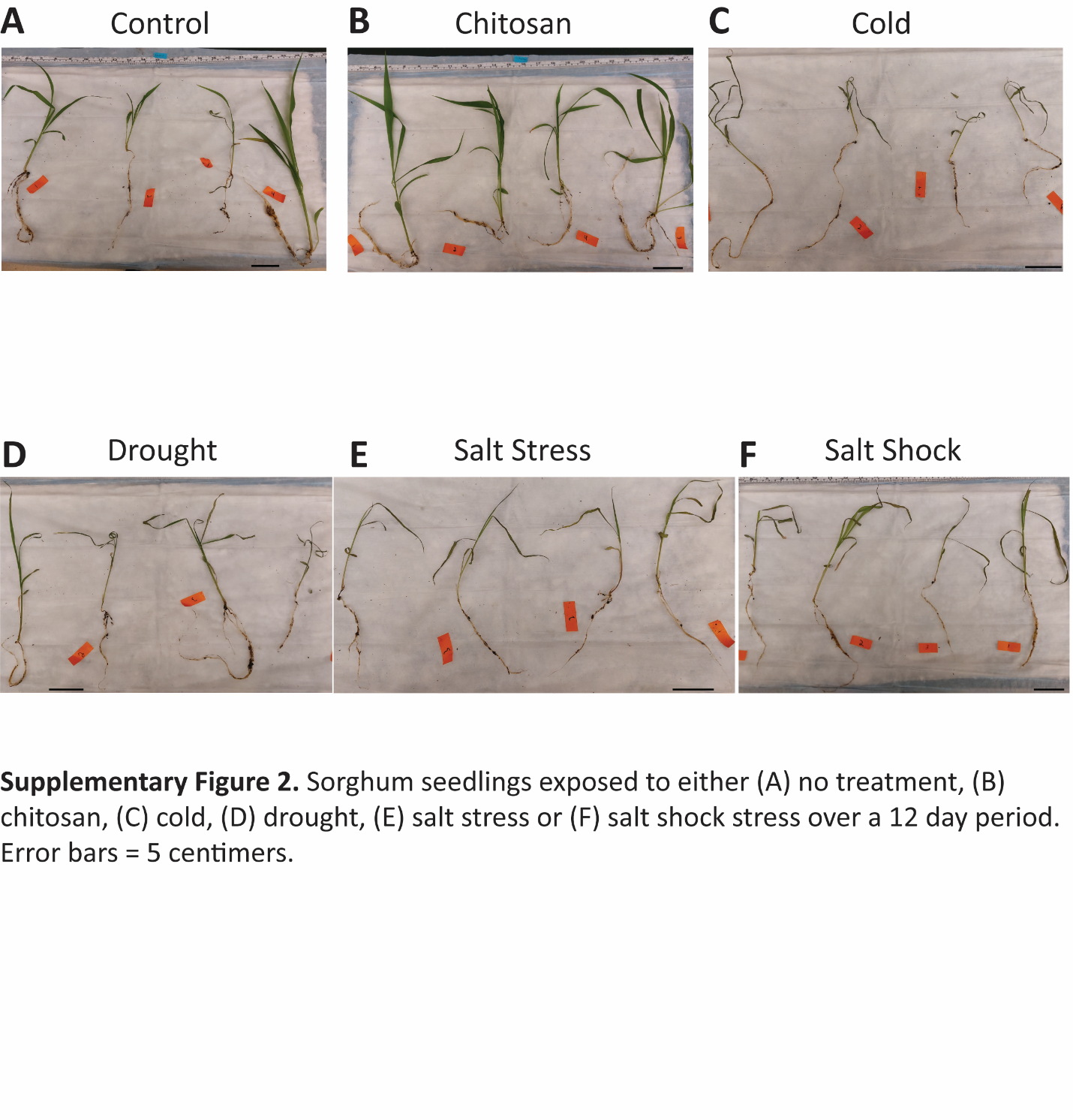

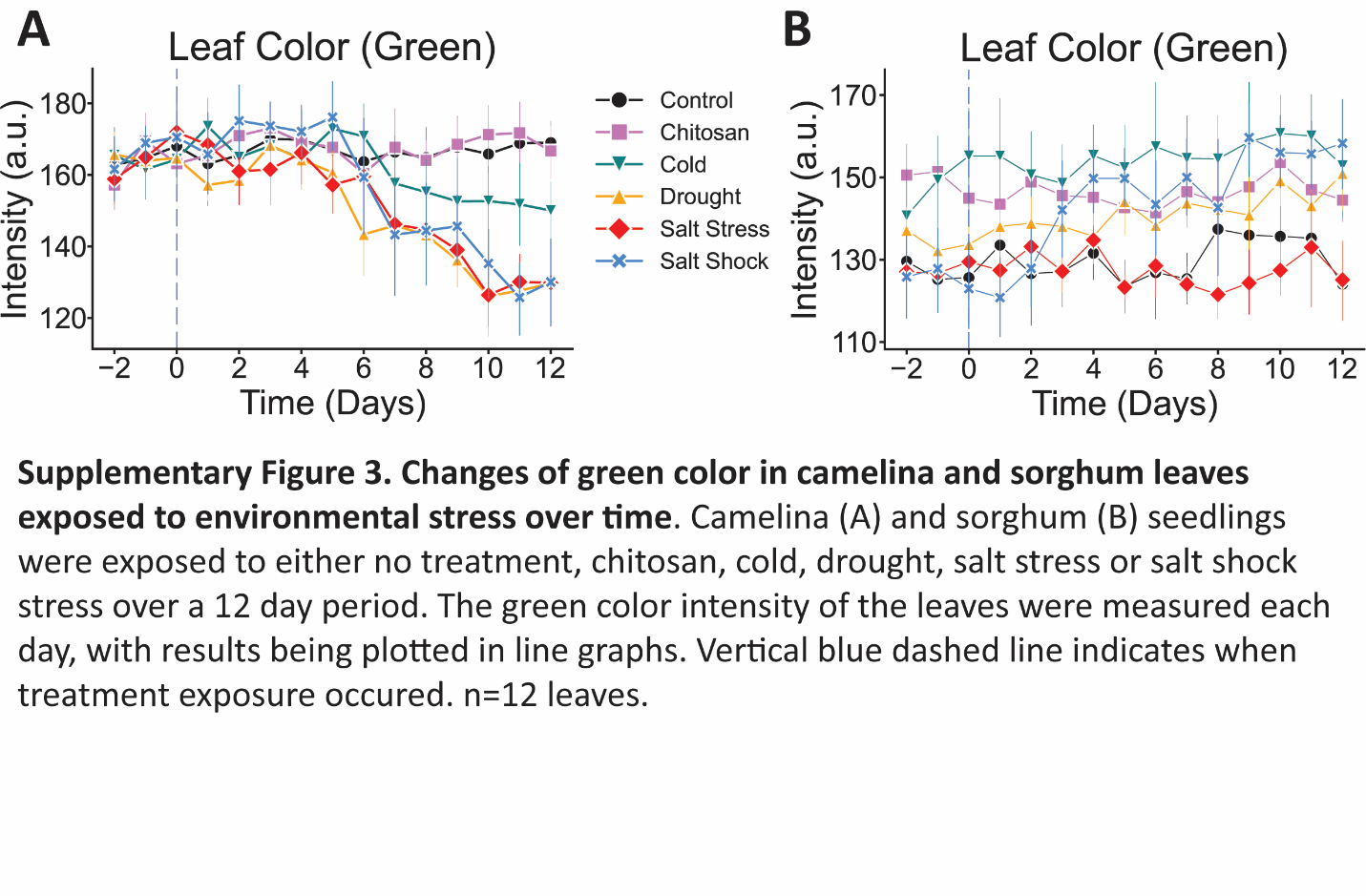

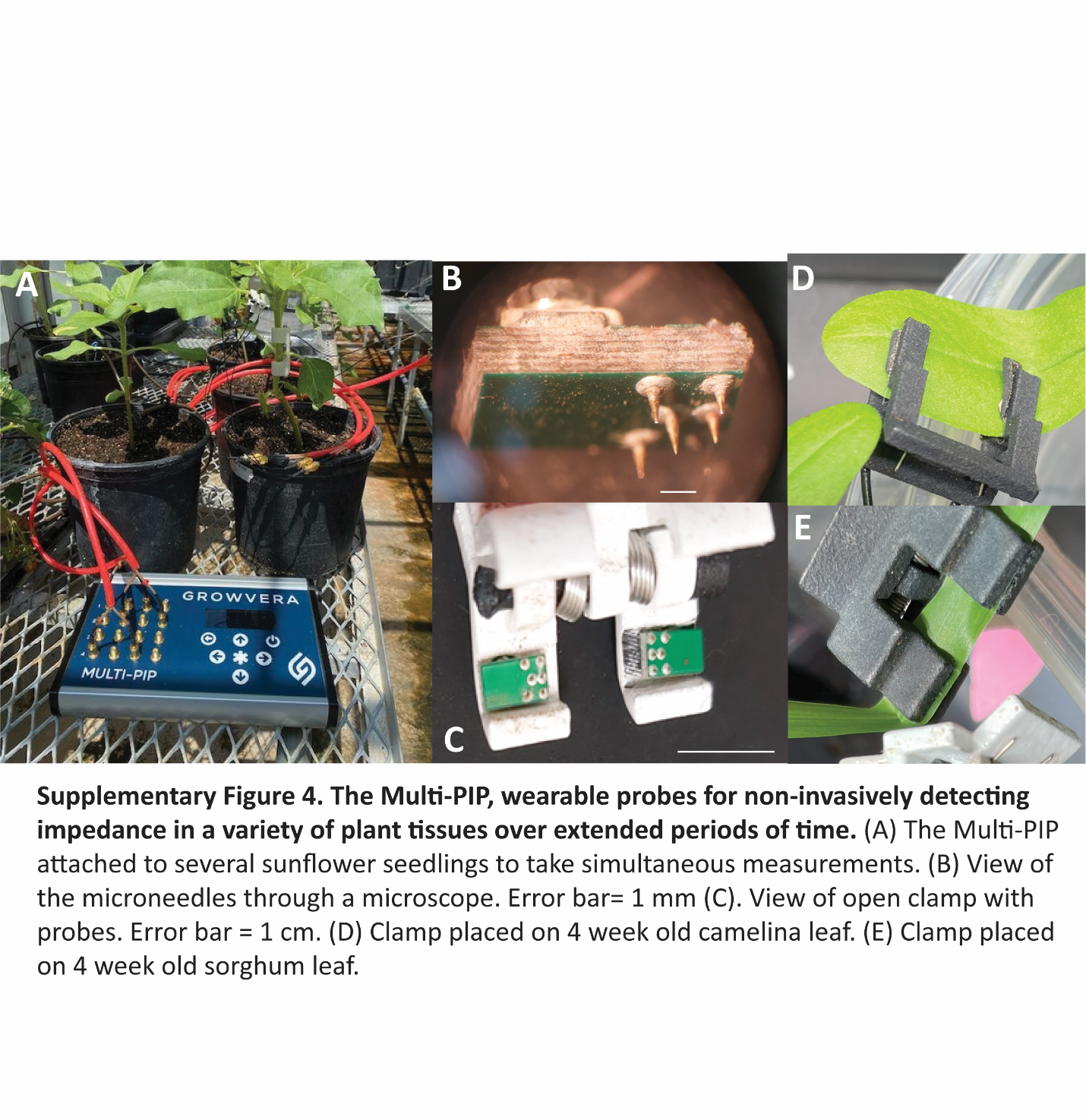

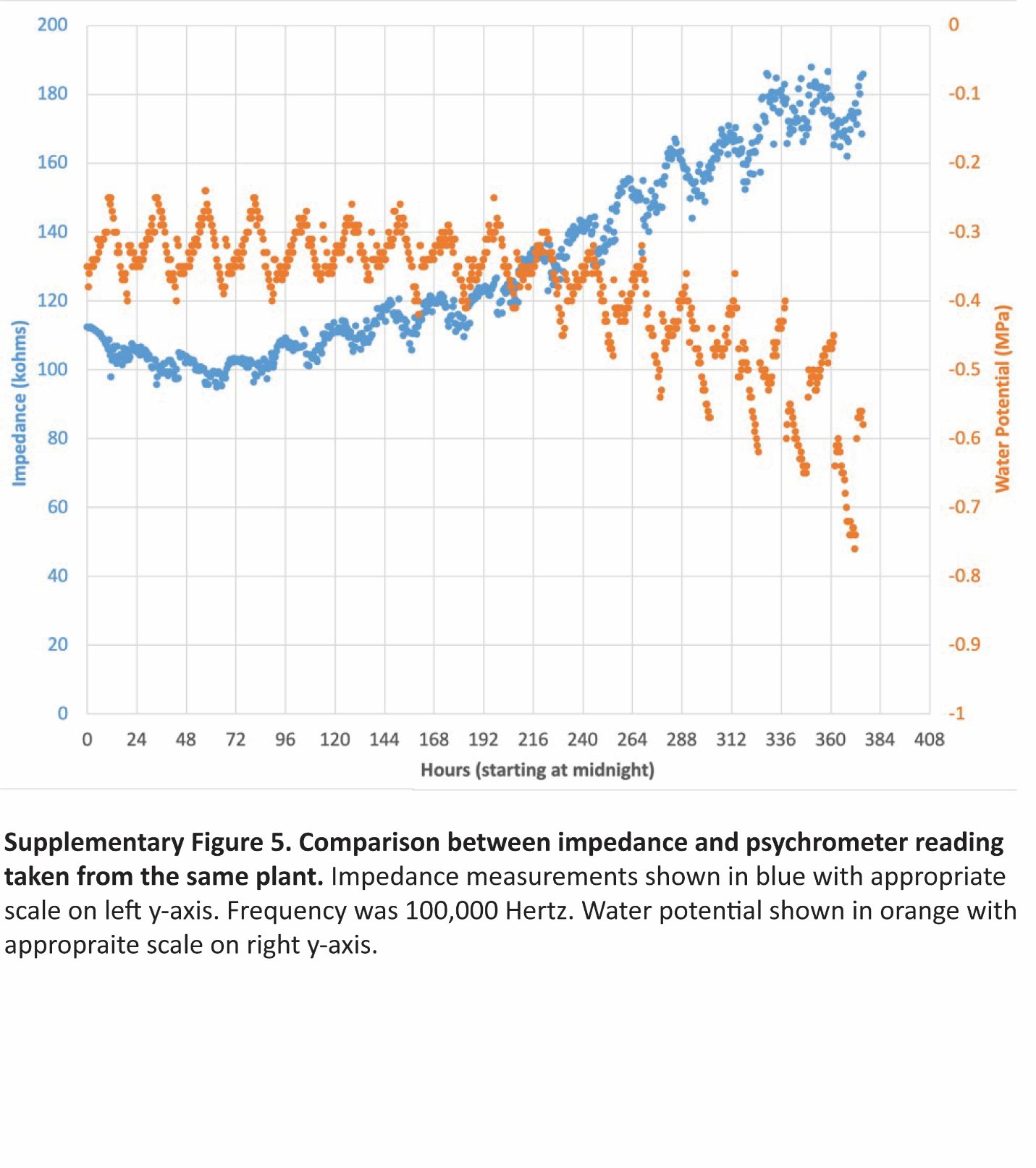

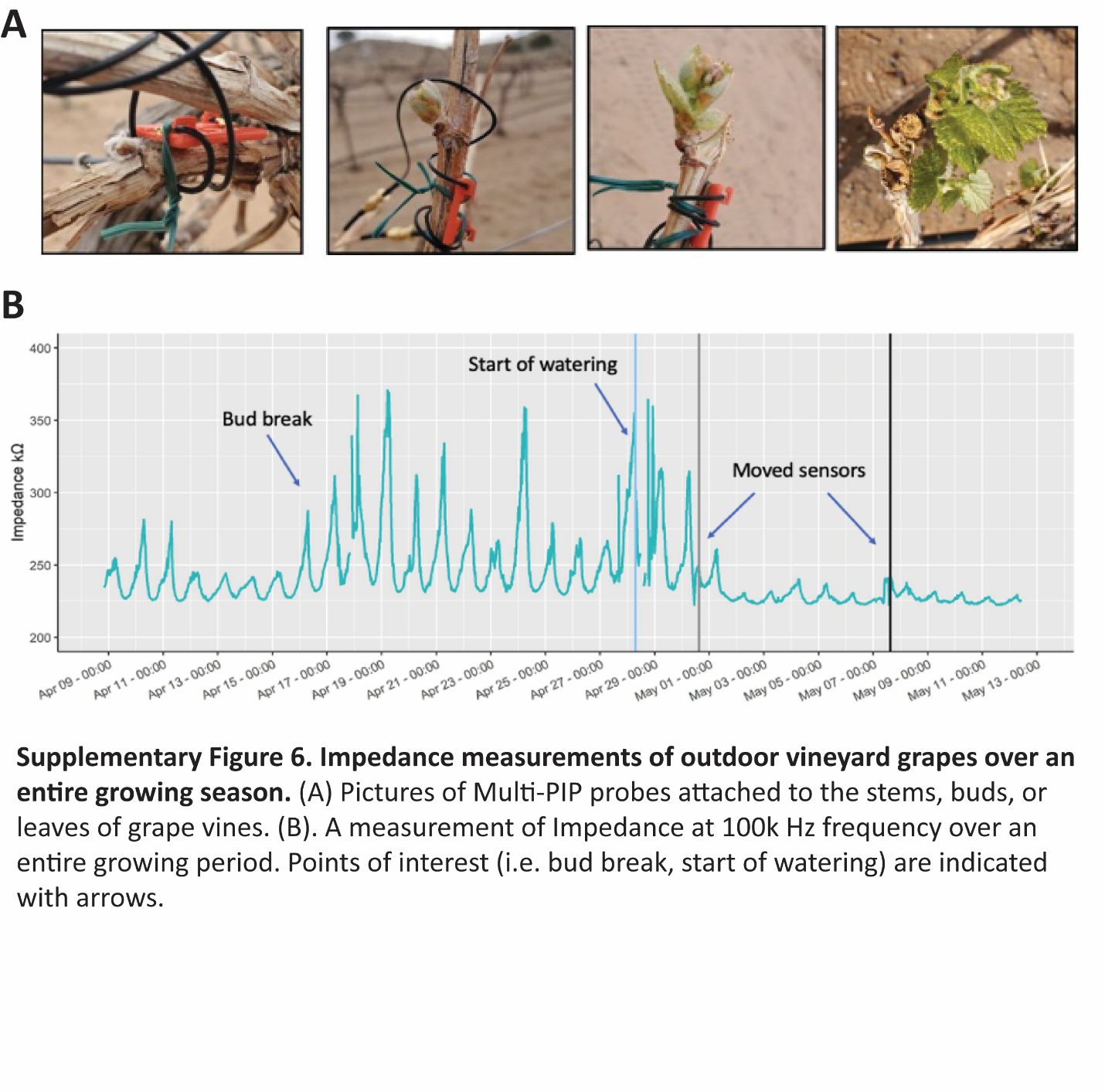

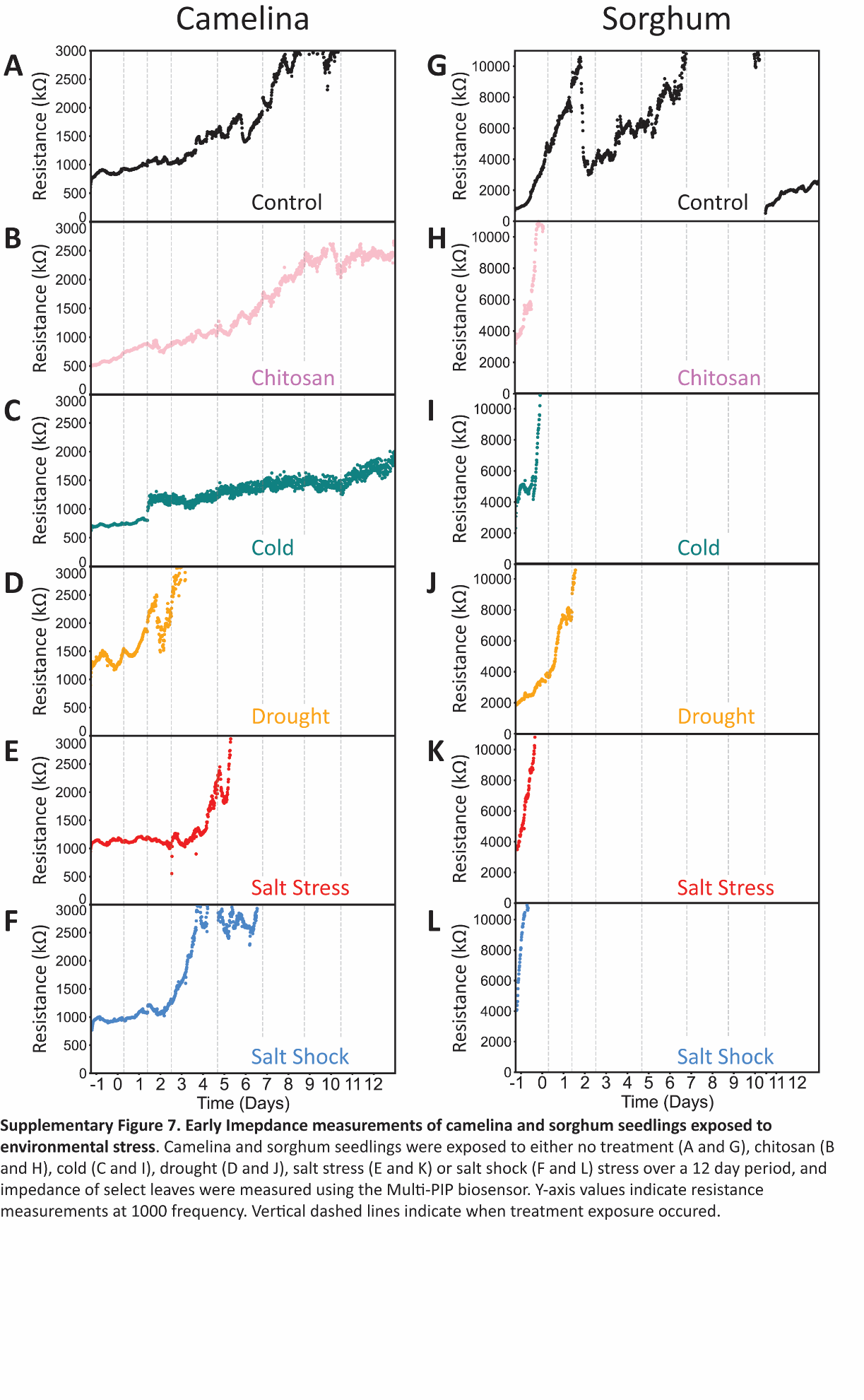

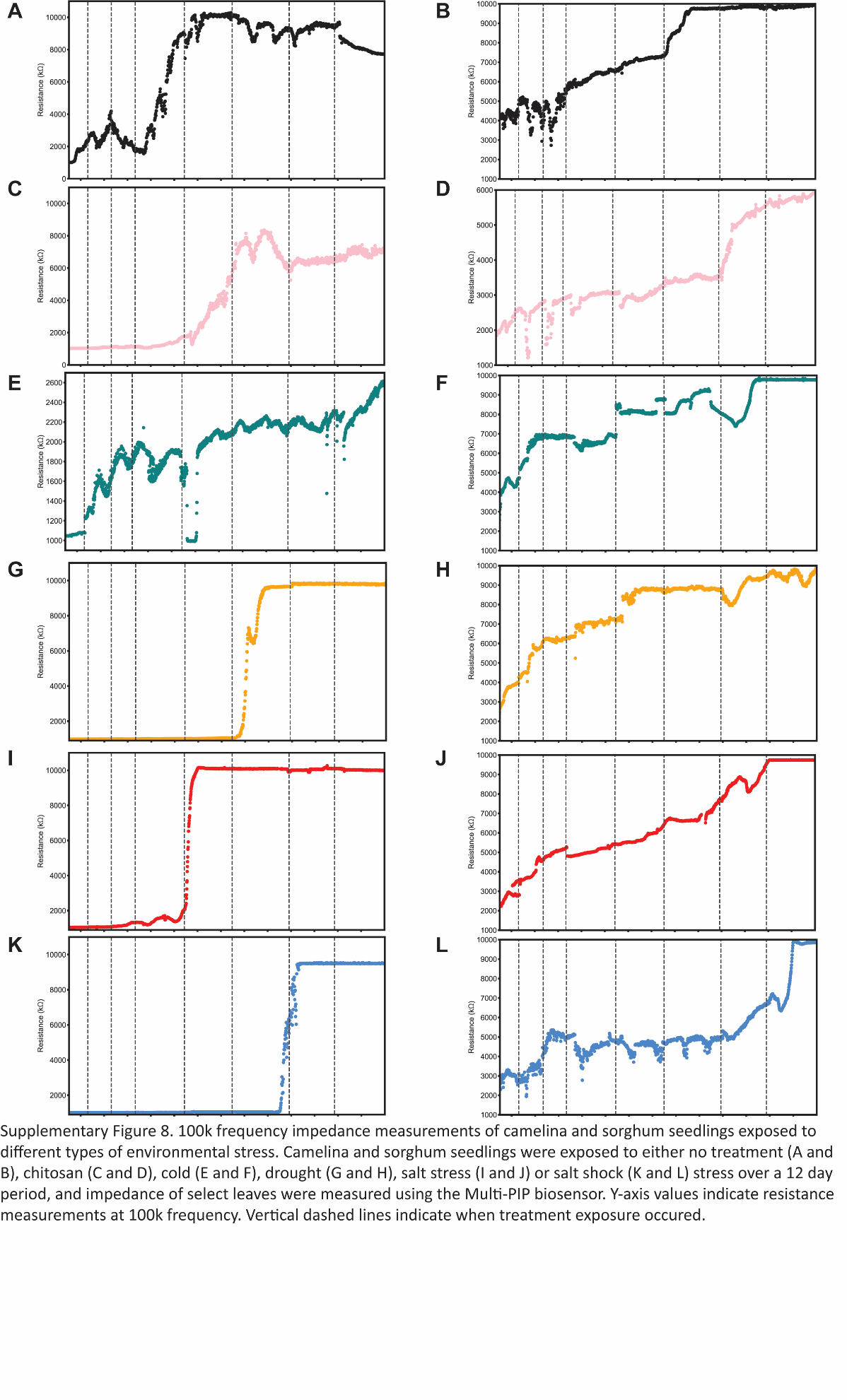

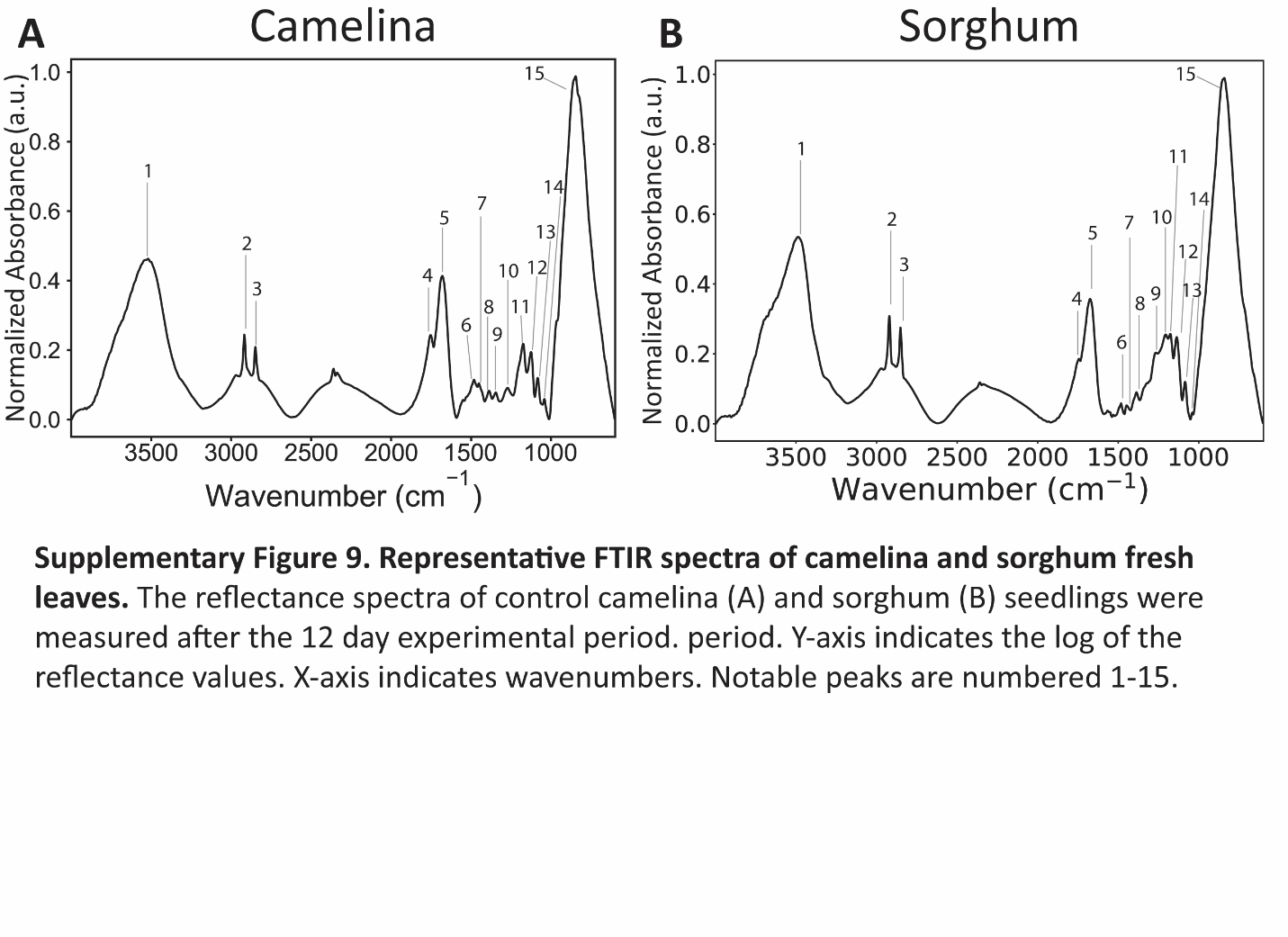

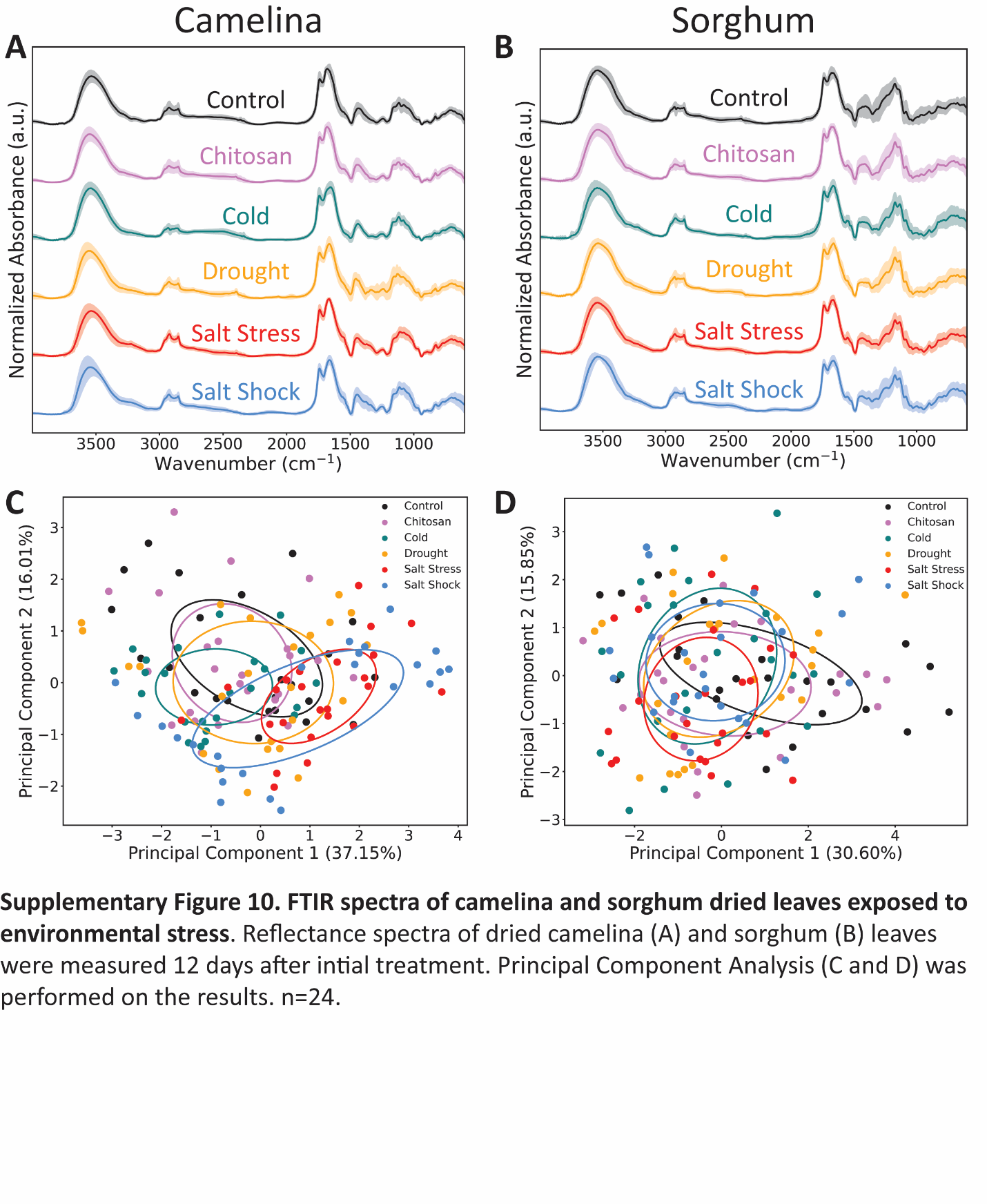

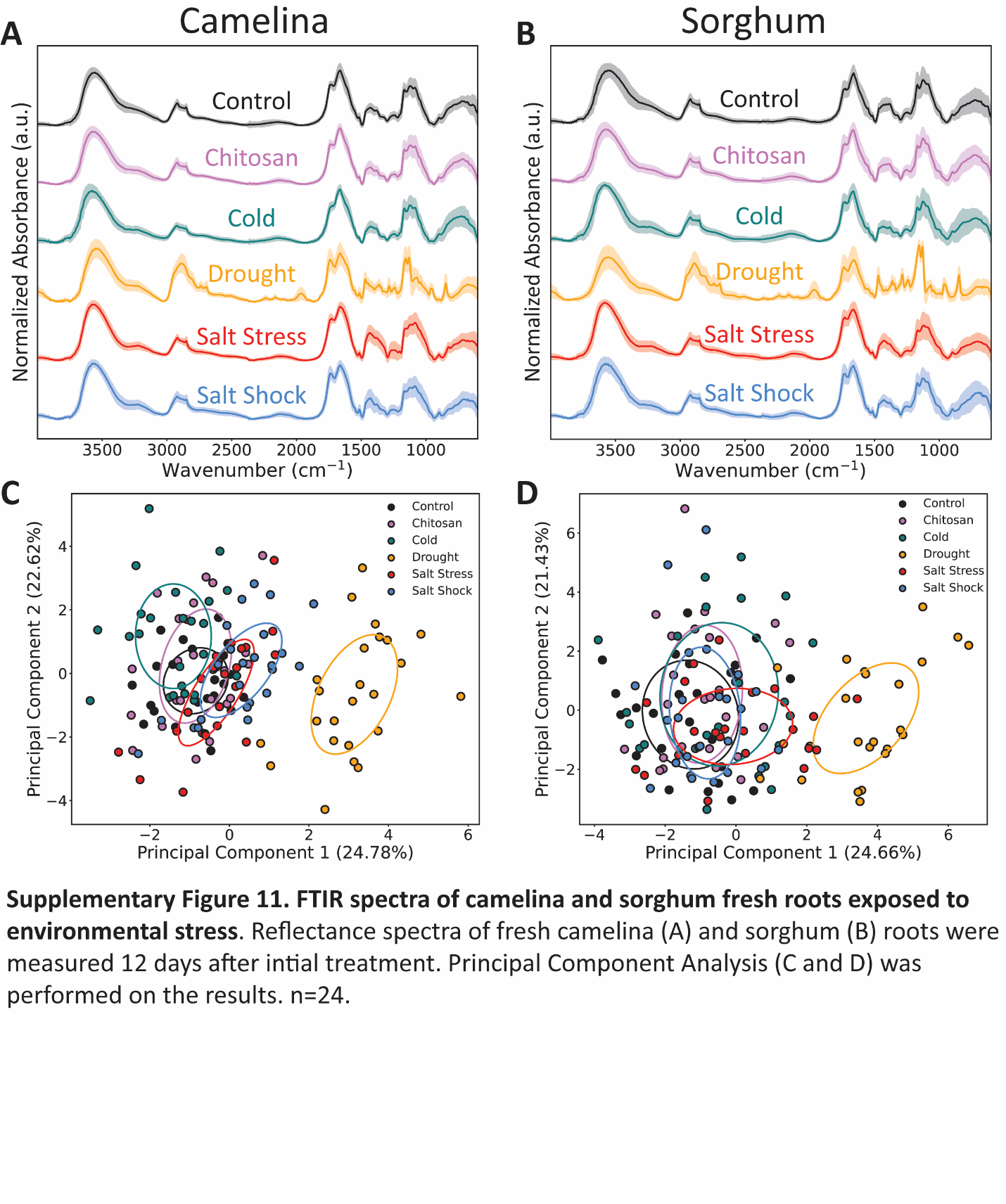

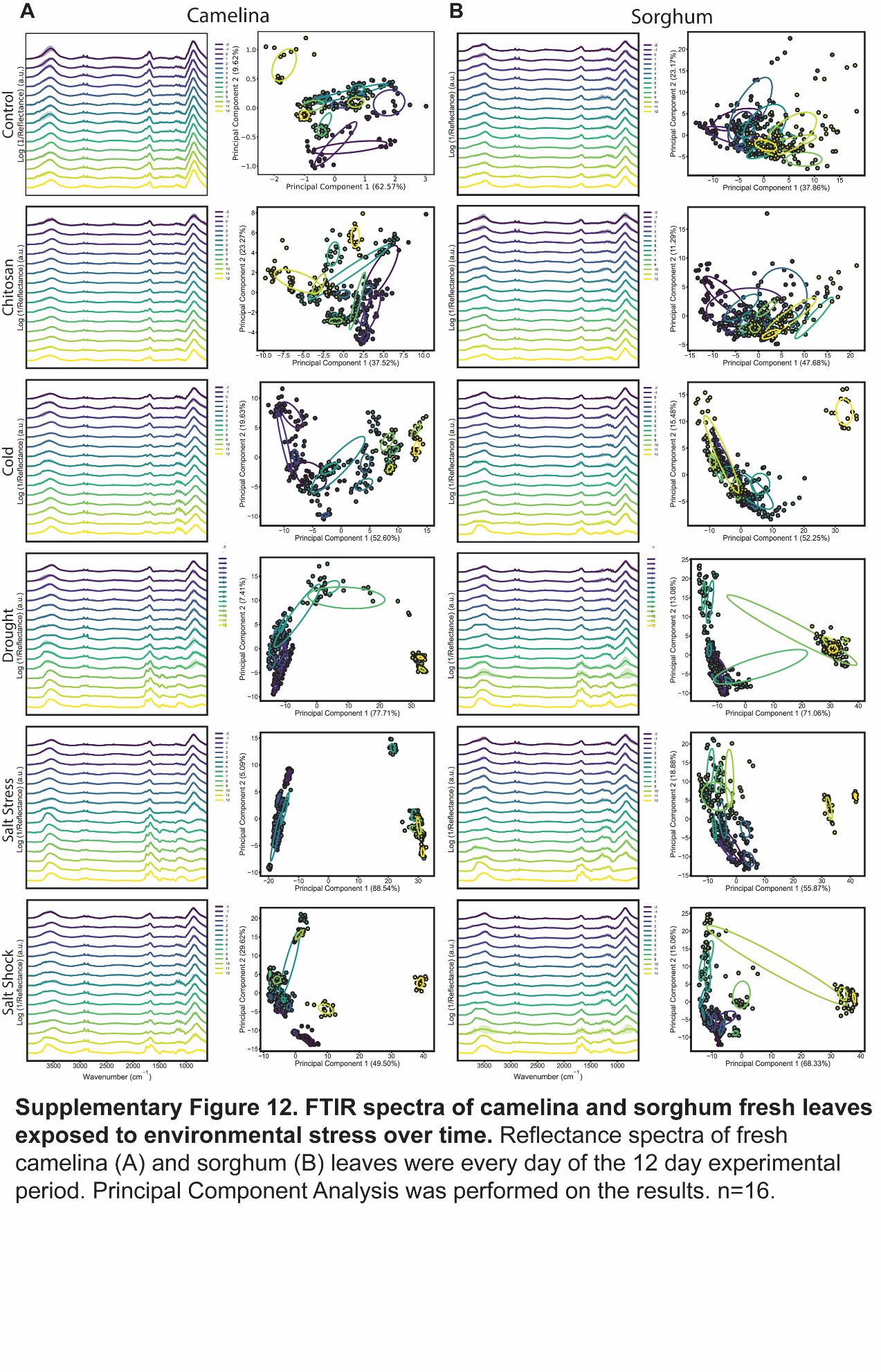

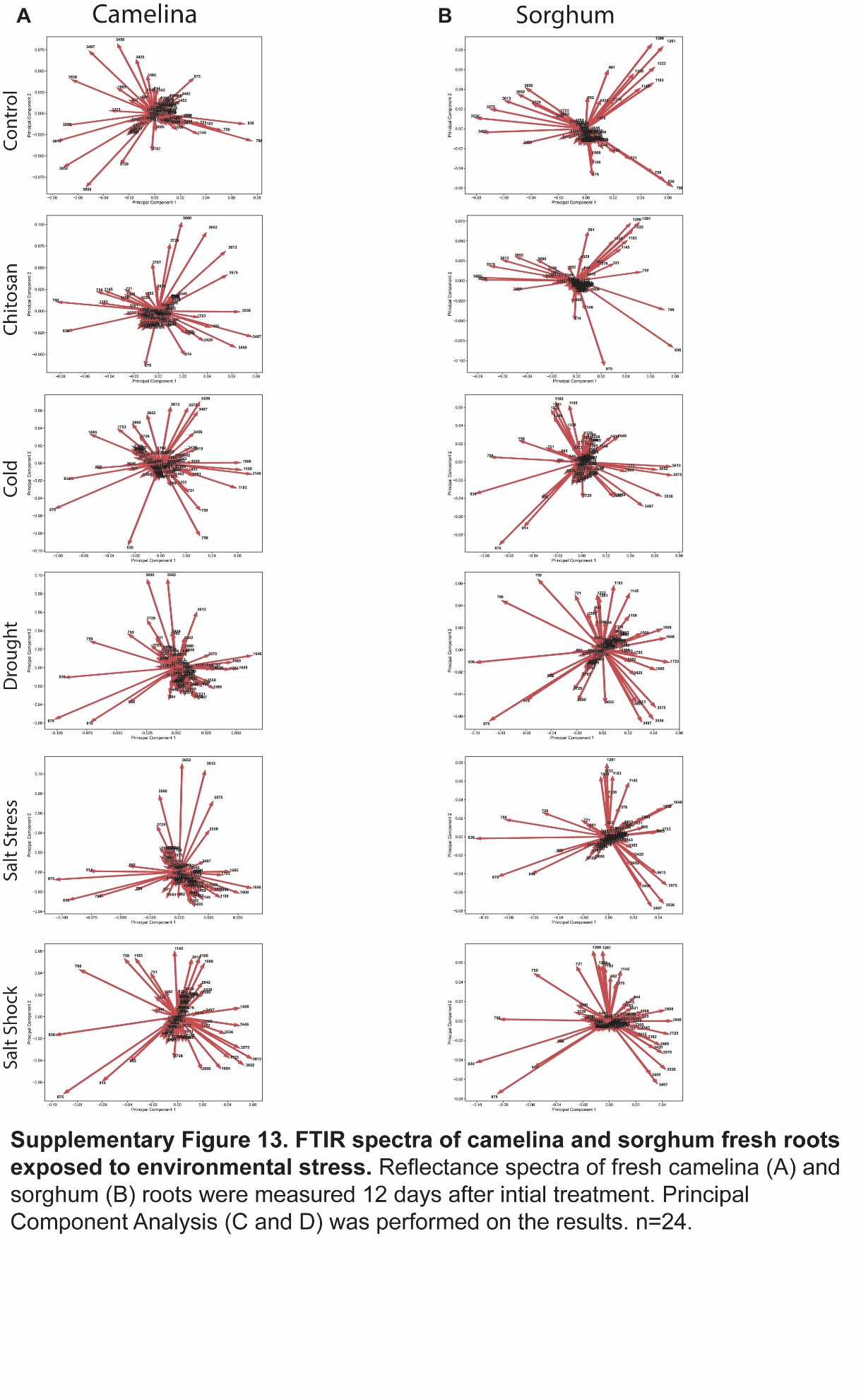

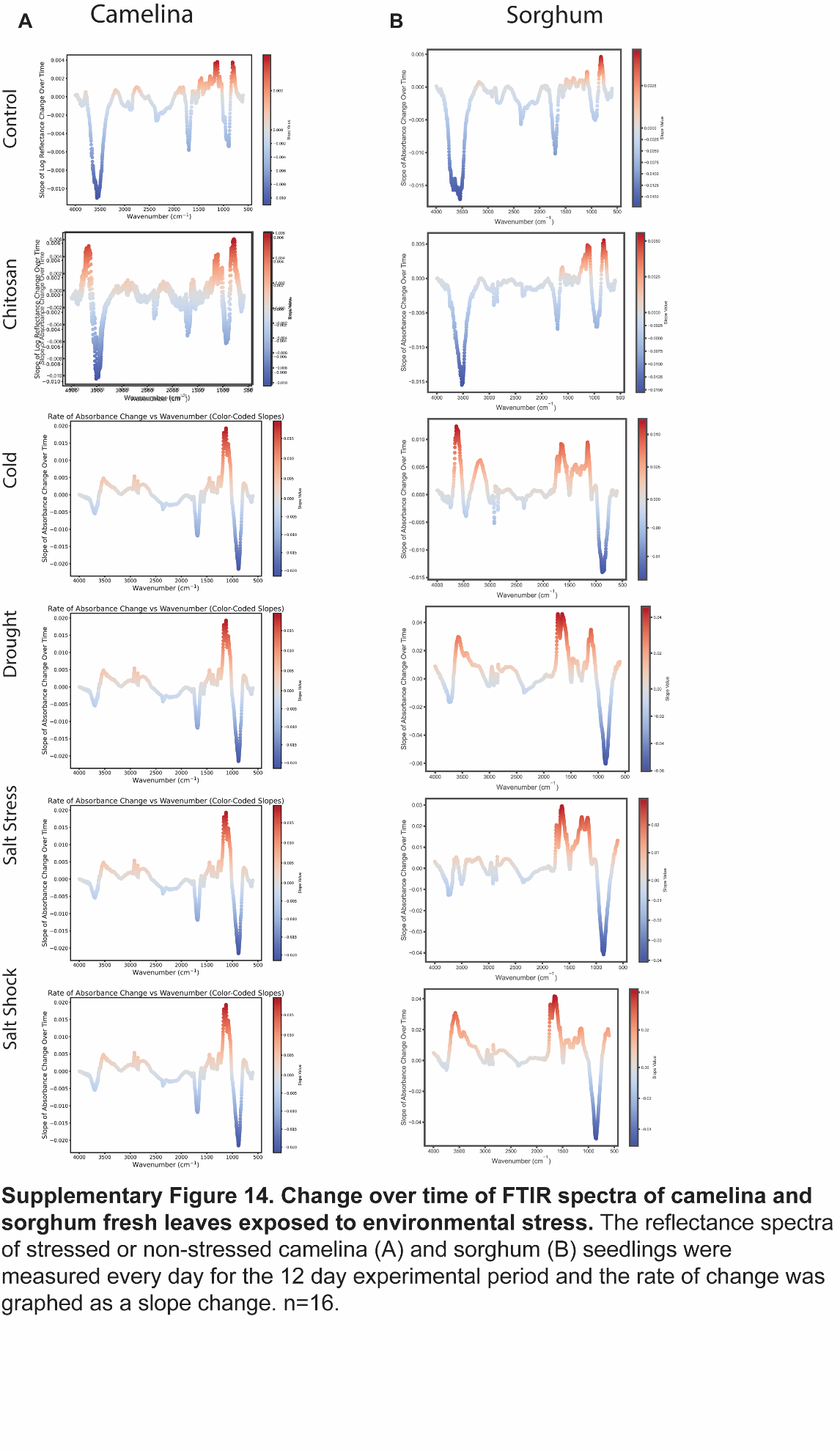

Table 3: Spectral Assignment of Peaks from Sorghum and Camelina Representative Reflectance Spectra.

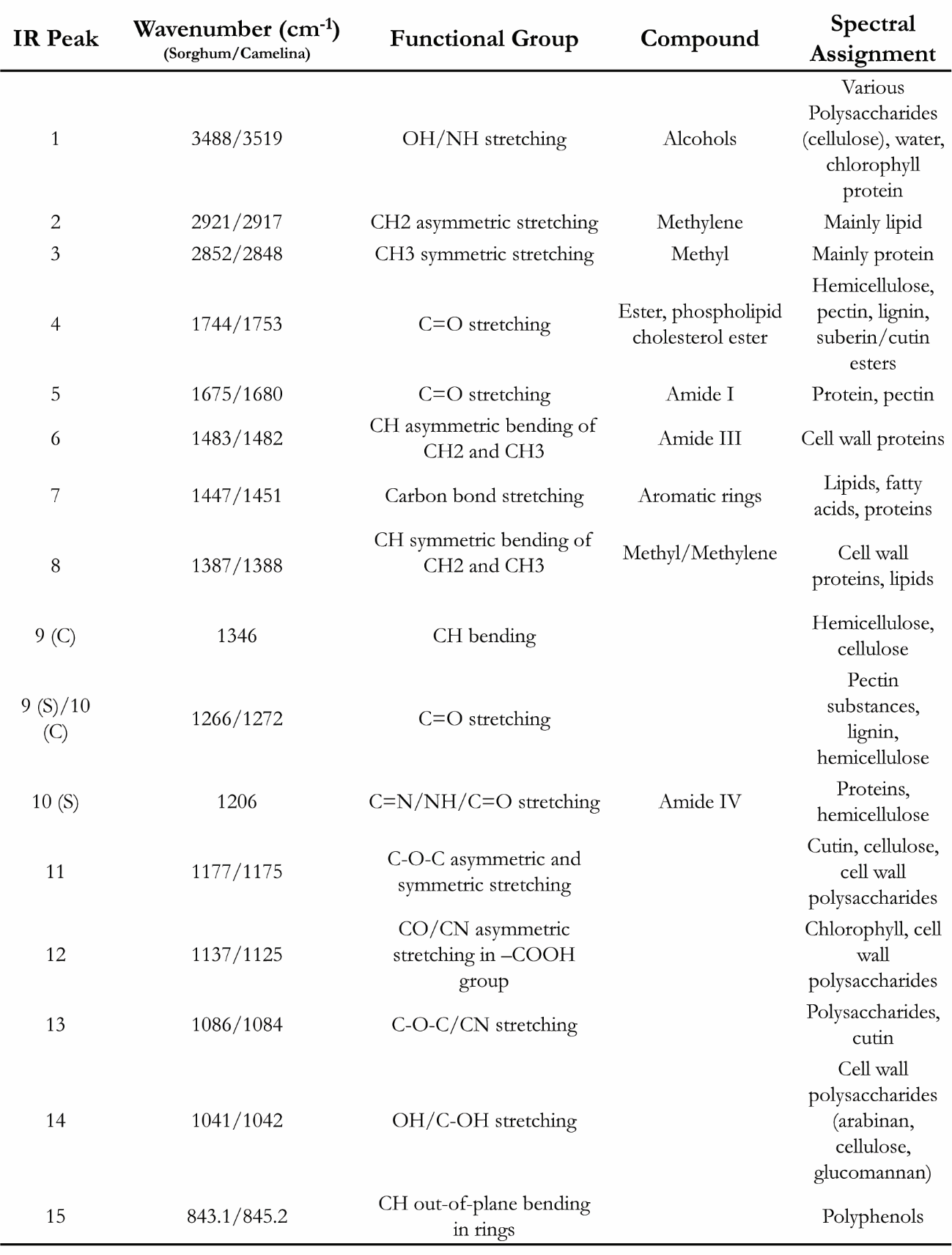
